# Diet Potentially Drives the Differentiation of Eating Behaviours via Alterations to the Gut Microbiome in Infants

**DOI:** 10.1101/2021.04.24.438478

**Authors:** Cathy Yan, Helen Zhao, Navika Nayar, Kyung E. Rhee, Julie C. Lumeng

## Abstract

Certain infant eating behaviours are associated with adverse health outcomes such as obesity. While a diet consisting of infant formula has been linked to higher-risk eating behaviours and changes in the gut microbiome, little is known about what role the gut microbiome plays in mediating eating behaviours. Using 16S rRNA sequences extracted from 96 fecal samples collected from 58 infants, we identified a subset of bacterial taxa that were more abundant in formula-fed infants, primarily composed of the phylum Firmicutes. The presence of these taxa correlated with a lower drive to eat (i.e., lower food responsiveness). Furthermore, short-chain fatty acid production pathways were significantly more abundant in formula-fed infants, negatively correlated with food responsiveness, and positively associated with relative abundance of the Firmicutes subset. Our results suggest that higher abundances of Firmicutes in formula-fed infants may decrease their food responsiveness through short-chain fatty acid production in the first four months of life. Taken together, these findings suggest a potential role for the infant’s diet in impacting eating behaviour via changes to the gut microbiome, which may lead to the development of novel interventions for the prevention of childhood obesity.

## INTRODUCTION

The human gut microbiome comprises trillions of different bacteria that interact to influence an individual’s physiology and mental health via immunologic, endocrine, and neural pathways (1). It can protect an individual by barricading pathogenic organisms from colonizing the body, aids in metabolism through processes that promote the breakdown of toxins or vitamin synthesis, and serves a trophic role by maintaining tolerance to antigens in food (1). For infants, their gut microbiota is similarly crucial for health and development, and is impacted by factors such as mode of delivery, exposure to antibiotics or probiotics, and diet (2).

Regarding diet specifically, previous literature has suggested that breastfeeding shapes the gut microbiota in neonates through direct introduction of the mother’s milk microbiota and prebiotics such as human milk oligosaccharides (HMO) (3). The consensus is that breastfed infants have gut microbiota with lower diversity and lower levels of the phylum Firmicutes compared to formula-fed infants (1, 4). Despite the impact of infants’ diet on specific taxa in their gut microbiota being explored, research regarding the effects of breastfeeding or formula-feeding on inter-microbial communities is lacking. As bacteria exist in complex networks rather than independently, this level of understanding is critical.

Additionally, breastfeeding may be associated with lower rates of childhood obesity through the theorized mechanisms of developing healthier food preferences and eating behaviours (5, 6). In infants, eating behaviours can be evaluated through the Baby Eating Behaviour Questionnaire, which measures several different eating behavior profiles: food responsiveness - the extent to which a child indicates an interest in and desires to spend time eating food; enjoyment of food - the extent to which a child finds eating pleasurable and desires to eat; satiety responsiveness - the extent to which a child becomes full easily and leaves food when finished eating; slowness of eating - the pace at which the child consumes their food; and general appetite, which correlates with the other metrics (7, 8).

An growing body of research indicates that an adult’s gut microbial profile may play a key role in their eating behaviours (9). A recent clinical study conducted by Sanmiguel et al. showed that interventions shaping the microbiomes of obese patients led to a reduction in their cravings (9). Other studies have shown that taking probiotic supplements decreases food intake in mice (10), and leads to weight loss in humans (11, 12). This may be because gastrointestinal microbes are incentivized to manipulate their hosts’ eating behaviour in order to minimize selective pressures, either by inducing intake of foods that “suppress their competitors”, or that “enhance their own fitness” (13). Furthermore, although previous studies support the gut-brain axis model where nervous stimulation by gut bacterial peptides results in activating the vagus nerve to regulate eating behaviours and body weight (14, 15), there remains a lack of research on how bacterial populations are associated with eating behaviours in infants.

In this study, we explored relationships between the infant’s diet, gut microbiome, and eating behaviours using the “eating behaviour development in infants” data repository by Rhee et al. Our aim was to examine how different diets influence the diversity and community composition of infants’ gut microbiomes, and explore how these microbiomes may relate to infant eating behaviors. Overall, we seek to propose a possible pathway for how the gut microbiota may influence eating behaviours, which could have implications for infants’ health. We hypothesize that the infant’s diet influences their gut microbial profiles, which, in turn, affects their eating behaviours. More precisely, we predict that formula-feeding is associated with a higher abundance of the phylum Firmicutes and the exhibition of obesity-prone eating behaviours.

## METHODS

### Participant Recruitment

Infant-mother dyads were recruited from the community. Mothers provided written informed consent for themselves and their infants. The University of Michigan Institutional Review Board approved this study. Inclusion criteria were: (1) Child was born at 37.0 – 42.0 weeks gestation, with weight appropriate for gestational age, and no significant perinatal or neonatal complications. Exclusions were: (1) non-fluency in English in the parent; (2) foster child; (3) mother < 18 years old; (4) medical problems or known diagnosis affecting current or future eating, growth or development; (5) child protective services involvement in the neonatal period; (6) infant does not consume at least 2 ounces in one feeding from an artificial nipple and bottle at least once per week. The exclusion of infants who had not yet taken a feeding from an artificial nipple resulted in the exclusion of few infants, as most infants in the population from which this cohort was recruited had fed from a bottle with an artificial nipple at least occasionally by the age of recruitment.

### Data Collection

At each age point, mothers completed the Baby Eating Behavior Questionnaire, an adapted version of the Children’s Eating Behaviour Questionnaire. The BEBQ is an 18-item parent-report psychometric measure of infant appetite. All questions in each subscale were scored on a 5-point Likert scale as never (1), rarely (2), sometimes (3), often (4), or always (5), and mean scores for each subscale were then calculated (range: 1–5). It generates 4 subscales (an example of a question is provided in brackets): Enjoyment of food (4 items; “My baby seemed contented while feeding”), Food responsiveness (6 items; Even when my baby had just eaten well s/he was happy to feed again if offered”), Slowness in eating (4 items; “My baby took more than 30 minutes to finish feeding”), Satiety responsiveness (3 items; “My baby got full before taking all the milk I thought s/he should have”), and General appetite (1 item). Internal reliability of the subscales by age based on Cronbach’s alpha were: Enjoyment of food (2 weeks: 0.62, 2 months: 0.61, 4 months: 0.70), Food responsiveness (2 weeks: 0.75, 2 months: 0.75, 4 months: 0.78), Slowness in eating (2 weeks: 0.63, 2 months: 0.62, 4 months: 0.57), Satiety responsiveness (2 weeks: 0.14, 2 months: 0.42, 4 months: 0.44). Given the poor internal reliability of the Satiety Responsiveness subscale in this sample, analyses of this subscale were omitted.

To assess infant dietary intake, we used selected questions from age-appropriate questionnaires developed by the U.S. Center for Disease Control (CDC); at each age point, mothers reported, in the last 7 days, the number of feedings per day of formula or breastmilk. From these data, infants were classified as exclusively breastfed, exclusively formula fed, or mixed. Only infants from the first two groups were included in subsequent analyses. At each age point, the mother reported, from a list of possible signs and symptoms (e.g., diarrhea, fever, vomiting), whether the infant had any health issues in the preceding two weeks. Mothers also reported whether they or their infant had taken any probiotics or antibiotics in the last two weeks. Mothers reported mode of delivery (Caesarean versus vaginal).

Fecal samples were collected from mothers and infants at each age point. Fecal samples from the initial cohort were collected using BD Swube™ dual headed. DNA was extracted and the 16S rRNA region was sequenced on an Illumina MiSeq platform using the 515F/806R primer set. Sequencing data was deposited in the European Nucleotide Archive (ENA) at EMBL-EBI under the accession number PRJEB39437 by the University of California San Diego Microbiome Initiative, with all other data recorded in the metadata.

### Identification of Confounding Variables

Potential factors that could influence the infant gut microbiome independent of diet were assessed based on prior knowledge and included maternal and infant intake of probiotics and/or antibiotics in the 2 weeks prior to sample collection, and mode of delivery (vaginal vs Caesarean-section). Associations between these factors, and diet and eating behaviours were evaluated using Fisher’s test and Mann-Whitney U test. Their impact on the gut microbiome was assessed based on weighted UniFrac distance and permutational analysis of variance (PERMANOVA) using the vegan package (16).

### Microbiome Sequences Analysis

Unless stated otherwise, the following analyses were performed using QIIME2 (v2020.11) and its plugins (17). Exact commands can be found in the supplementary command line script. After demultiplexing, 16S rRNA sequences underwent quality control using DADA2 (18). Next, a phylogenetic tree was generated and used to plot an alpha-rarefaction curve to identify the sampling depth at which richness has been fully observed. Taxonomy was assigned using the Greengenes 99% OTU database (19).

### Alpha and Beta Diversity Calculations

Metadata, along with the phylogenetic tree and taxonomy-annotated feature table exported from QIIME2, were imported into R. *Ape* (20) was used to convert QIIME2’s multichotomous tree into a dichotomous one for downstream analyses. *Phyloseq* (21) and *btools* were used to calculate alpha and beta diversity metrics for comparing breastfed and formula-fed infants. *Phyloseq* was used to calculate observed OTUs, Chao1, ACE, Shannon, Simpson, Inverse Simpson, and Fisher alpha diversity metrics, and Bray-Curtis, Jaccard, weighted UniFrac, and unweighted UniFrac distances for beta diversity. *Btools* was used to calculate Faith’s phylogenetic diversity. Statistical significance was evaluated using the Mann-Whitney U test for alpha diversity and PERMANOVA for beta diversity.

### Random Forest Classifier

Using *caret* (22) and *randomForest* (23), a random forest classifier was optimized, trained, and used to predict diet based on genus-level relative abundance. Receiver operating characteristic curves and feature importance were also calculated using these two packages.

### Co-abundant Clusters Identification

Microbial co-abundance at the genus level was calculated for genera that were present in at least twenty percent of the infants. Spearman correlation distance and Ward’s linkage were calculated for the centre log ratio-transformed relative abundance values and used to cluster microbes, as previously described by Cirstea et. al (24). The Mann-Whitney U test was used to compare relative cluster abundance between breastfed and formula-fed infants. Spearman correlation was calculated to assess the correlation between cluster relative abundance and eating behaviours.

### Metabolic Pathways Analysis

Inferred functional microbiota profiling was done using PICRUSt2 (v2.3.0b) (25). Differences in the relative abundance of metabolic pathways present in at least five percent of the infants were assessed using *ALDeX2* (26). Pathways were deemed to be statistically significantly differentially present when the Benjamini-Hochberg corrected P values for Welch’s t-test and the Wilcoxon test were both less than 0.05.

## RESULTS AND DISCUSSION

### Participant Characteristics

This study uses data collected from 58 infants at ages 2 weeks, 2 months, and 4 months for a total of 96 samples. These infants were exclusively breastfed or formula-fed in at least the 7 days leading up to data collection, and had no vomiting, diarrhea, or fever. Sample characteristics are shown in Table 1. Because the sample size did not provide sufficient power for a linear mixed effects (LME) model with infant ID as a random effect and age as a nested random effect, all samples were treated as independent even if they came from the same infant at different timepoints. Some infants provided only one sample, making a random-slope and random-intercept LME model infeasible. An LME model with only infant ID as the random effect indicated no significant associations between the gut microbiome and eating behaviours.

**Table 1.**
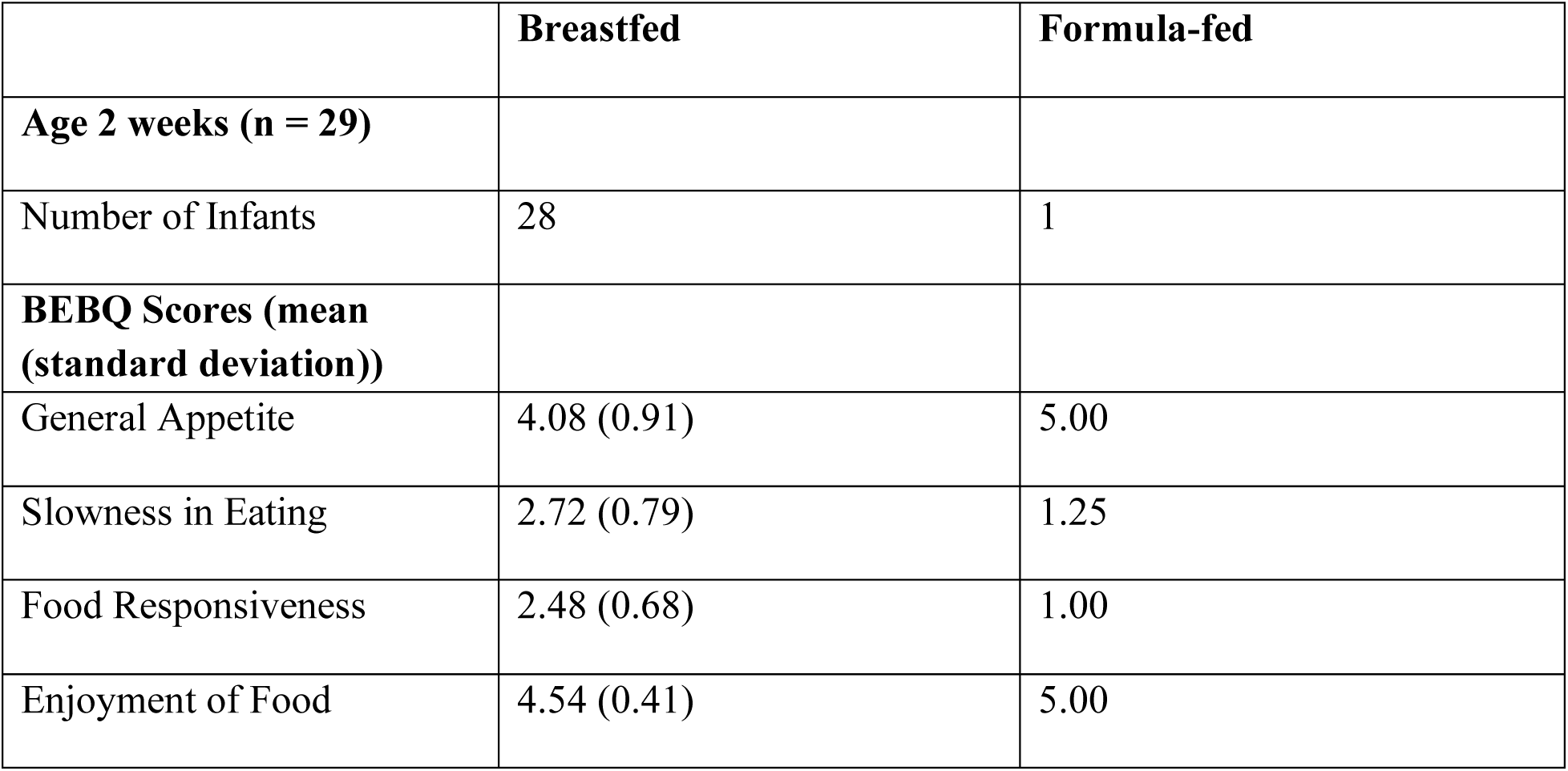
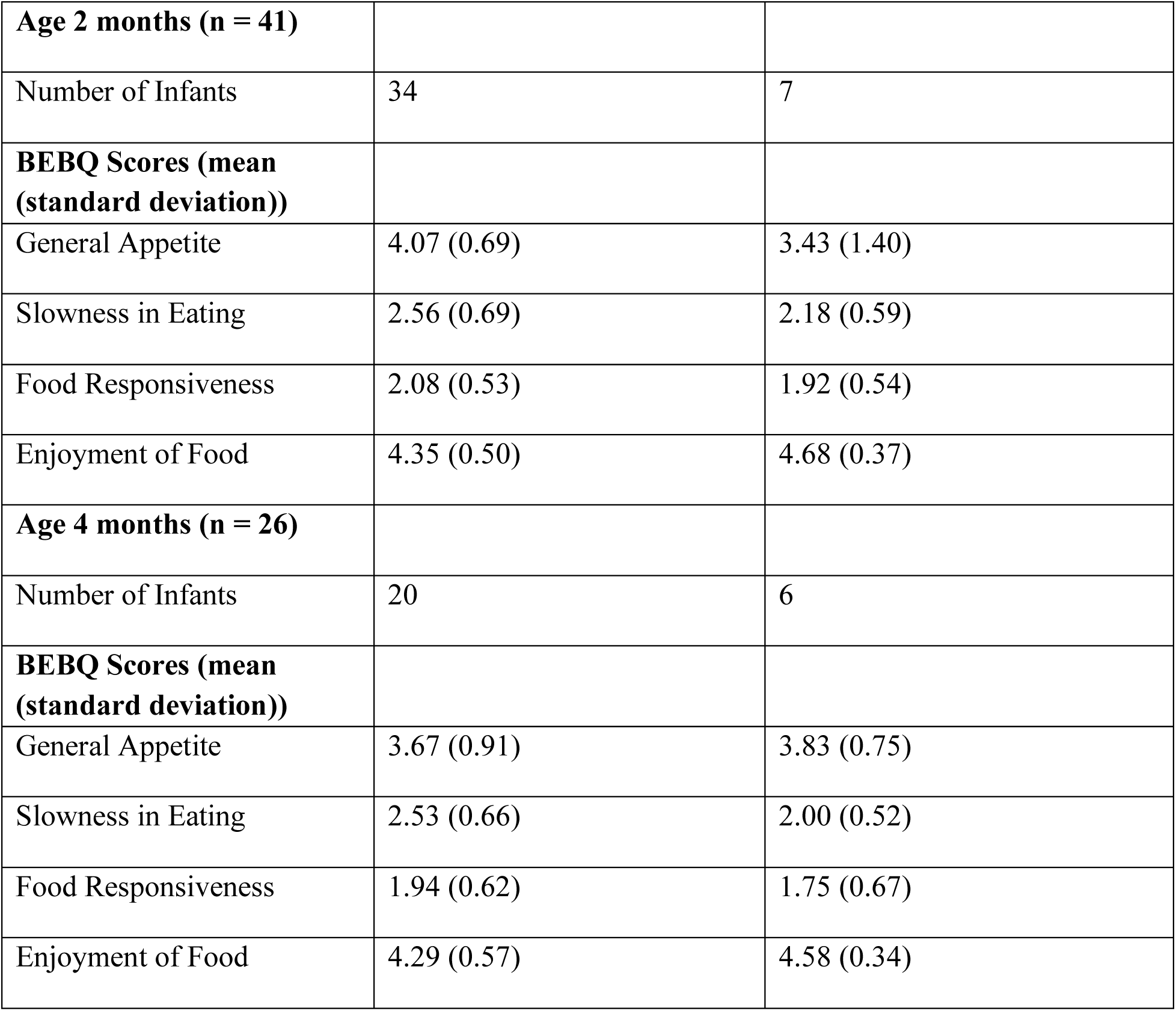
Sample Characteristics (n = 58). BEBQ scores are on a scale of 1 to 5, with higher scores indicating greater demonstration of the behaviour.

### No confounding variables were identified

In addition to diet, our main variable of interest, the mode of delivery (19), antibiotic use (20), and probiotic use (21) have been reported to impact the infant gut microbiome. Consequently, we assessed the effect of each factor within our study cohort. As the PCoA plot for weighted UniFrac distance accounted for the most variance compared to those based on Jaccard, Bray-Curtis, and unweighted UniFrac distances (Fig. 1a, Supplementary Fig. 1), we used weighted UniFrac as our metric for evaluating the impact of confounders on the gut microbiome. Additionally, we also ensured that no other confounding variable was associated with the infant’s diet. We found that infant gut microbiomes did not cluster differently based on weighted UniFrac distance by probiotic usage (PERMANOVA: F_95_ = 0.55, P = 0.67, R^2^ = 0.0059), antibiotic usage (PERMANOVA: F_95_ = 1.41, P = 0.22, R^2^ = 0.015), or mode of delivery (PERMANOVA: F_95_ = 1.67, P = 0.122, R^2^ = 0.035).

**FIG. 1.**
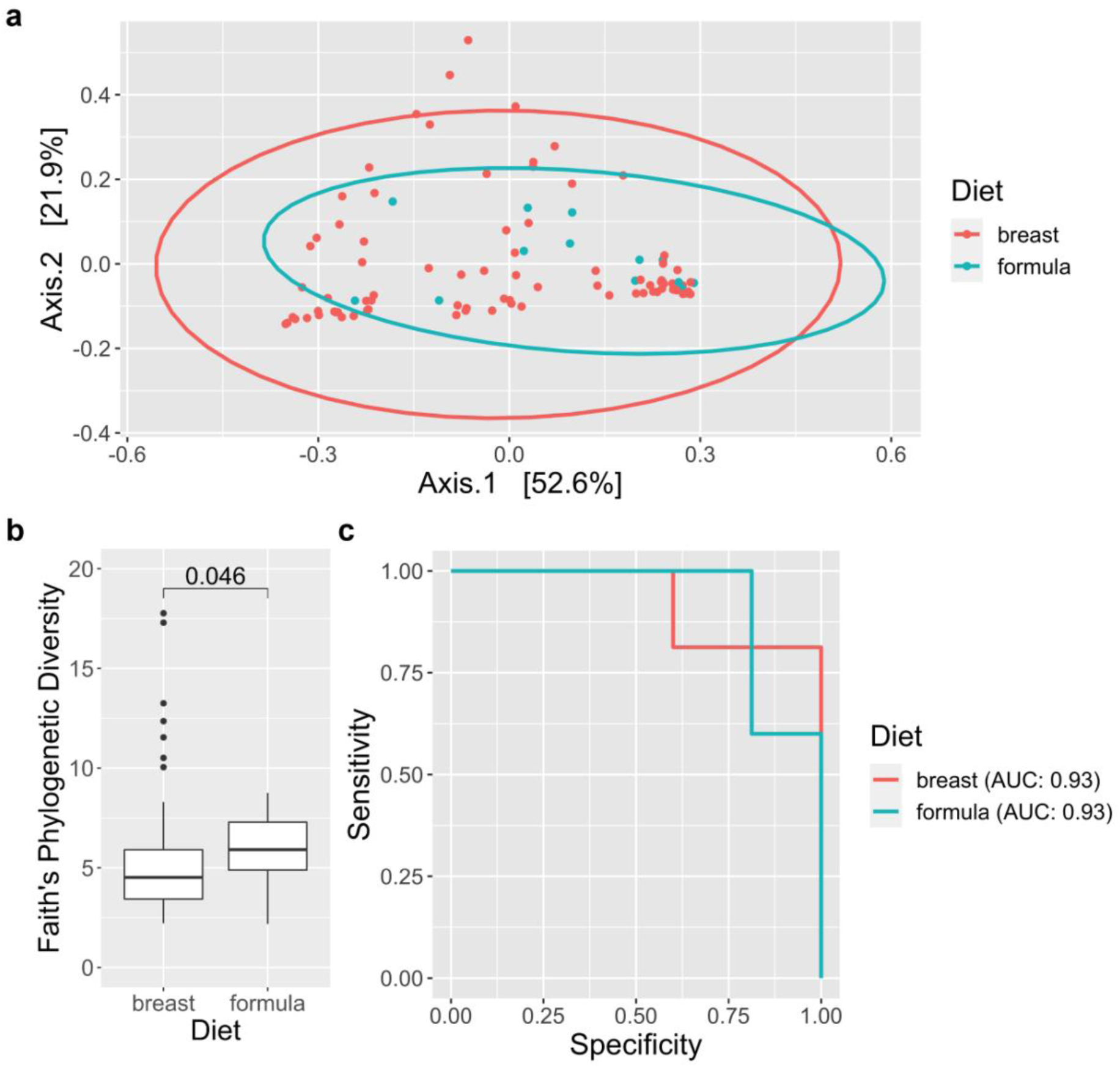
Breastfed and formula-fed infants host distinct microbiomes. (A) Weighted UniFrac beta diversity PCoA plot for infant samples, coloured by diet (breastfed or formula-fed). Statistical significance was assessed using PERMANOVA (F_95_ = 3.69, P = 0.02, R^2^ = 0.038). (B) Comparison of Faith’s phylogenetic diversity between diet. Whiskers represent 1.5 times the interquartile range; points beyond them represent outliers. Statistical significance was assessed using the Mann-Whitney U test. (C) Receiver operating characteristic (ROC) curve for evaluating the performance of a random forest classifier trained to separate breastfed and formula-fed infants.

### Breastfed and formula-fed infants host distinct microbiomes

We compared alpha and beta diversity metrics for breastfed and formula-fed infants, expecting the two groups to host distinct microbial communities and for breastfed infants to have lower alpha diversity (27). For these analyses, we started by determining the sampling depth at which an increase in depth led to no change in alpha diversity. Based on the alpha-rarefaction curve (Supplementary Fig. 2a) and reads frequency histogram (Supplementary Fig. 2c) generated using QIIME2, the sampling depth was set at 14,000 reads. This sampling depth led to 17 infants being excluded from diversity analyses.

Next, weighted UniFrac was again used as the beta diversity metric for how infants clustered based on diet (Fig. 1a). Although our PCoA displayed no obvious visible clustering, the gut microbiomes of breastfed and formula-fed infants were significantly different (PERMANOVA: F_95_ = 3.69, P = 0.02, R2 = 0.038). For alpha-diversity, formula-fed infants had higher levels of alpha diversity than breastfed infants across most metrics (Observed: P < 0.001, Chao1: P = 0.0011, ACE: P = 0.001, Shannon: P = 0.012, Simpson: P = 0.1, Inverse Simpson: P = 0.1, Fisher: P < 0.001); Supplementary Fig. 3). Breastfed infants also had significantly lower Faith’s phylogenetic diversity than formula-fed infants (P = 0.046; Fig. 1b).

The fact that breastfed and formula-fed infants differed in terms of weighted UniFrac and multiple alpha diversity metrics suggests differences in taxonomic composition between the two diets. However, instead of merely identifying differentially abundant taxa, we attempted to distinguish between breastfed and formula-fed infants based on relative abundance at the genus level using a random forest classifier. This strategy has been used previously to successfully separate healthy dogs from those with irritable bowel syndrome (28). With our model, we achieved an area under the curve (AUC) of 0.87 (Figure 1c). These data demonstrate that breastfed and formula-fed infants’ gut microbiomes are distinct in genus-level composition, particularly for a phylum reported to be more abundant in formula-fed infants (29).

### Two communities of bacteria are differentially abundant between breastfed and formula-fed infants

Since most of the genera that best distinguish between infants with different diets belonged to the phylum Firmicutes, we tested if those genera are related functionally. Within the gut, microbes are part of networks that cooperate and compete (30). We inferred the presence and composition of these types of communities based on genus-level coabundance. Spearman’s correlations between genera present in at least twenty percent of the infants were calculated and used for clustering into a dendrogram. Covariance was then visualized using a heatmap, and three clusters of high covariance composed of at least three genera were identified (Supplementary Figure 4a). Out of these, only two were found to be differentially abundant between breastfed and formula-fed infants (Supplementary Figure 4b; Figure 2a).

**FIG. 2.**
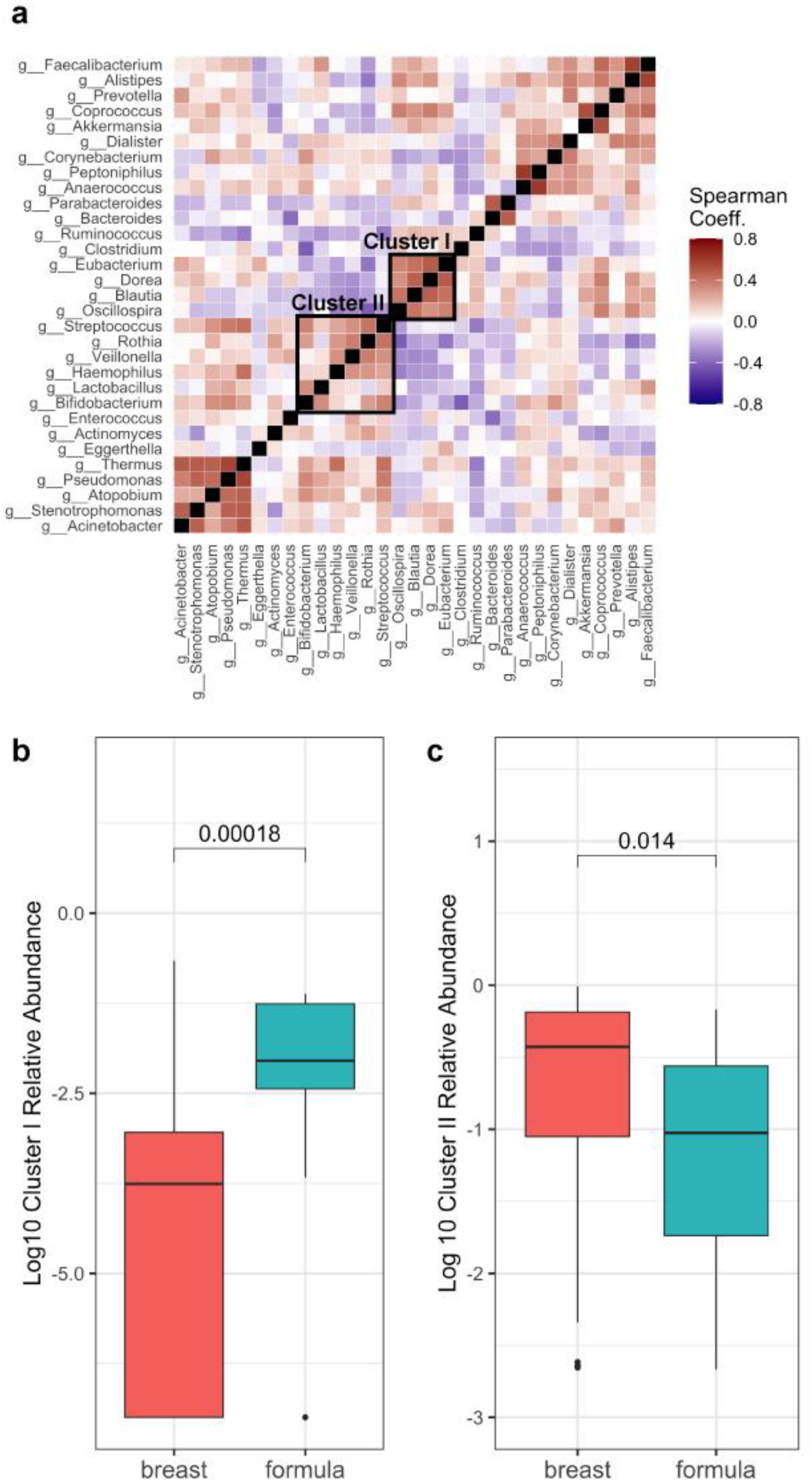
Breastfed and formula-fed infants host two microbial communities that are differentially abundant. (A) Heatmap based on the covariance of bacterial genera coloured by Spearman correlation coefficient. Axes are arranged based on Spearman correlation distance and Ward linkage. (B, C) Two covariant bacterial clusters are differentially abundant between breastfed and formula-fed infants based on the Mann-Whitney U test.

The first, Cluster I, is composed of the genera *Dorea, Eubacterium, Blautia*, and *Oscillospira*, and is significantly more abundant in formula-fed infants (P < 0.001; Figure 2b). This is in concordance with previous studies reporting decreases in levels of the phylum Firmicutes and order Clostridiales in breastfed infants (31). *Blautia* regulates G-protein coupled receptors through butyric and acetic acid production, decreasing obesity and visceral fat accumulation (32). *Oscillospira* is also associated with leanness as it degrades animal-derived glycans from the host (33). *Eubacterium* is a prolific butyrate producer, breaking down complex carbohydrates from dietary fibers (34).

Cluster II, composed of the genera *Bifidobacterium, Lactobacillus, Haemophilus, Rothia, Streptococcus*, and *Veillonella*, is significantly more abundant in breastfed infants (P = 0.014; Figure 2c). *Bifidobacterium* is well-known to dominate the microbiota of breastfed infants, with some studies reporting as much as double the relative abundance in breastfed infants compared to formula-fed infants (35). The same study also noted higher levels of *Lactobacillus* and *Streptococcus. Bifidobacterium* and *Lactobacillus* digest dietary fibers and produce acetate (36), while *Streptococcus* and *Veillonella* induce cytokine production to modulate the gut immune system (37). *Rothia* degrades gluten (38). Additionally, research has shown that genera in Cluster II make up a large proportion of the breast milk microbiota (39).

### Diet is a driver of the relationships between food responsiveness and relative cluster abundance

The composition of the gut microbiome has been found to impact eating behaviours, but most research has involved adult rather than infant cohorts, and focused on individual taxa rather than microbial communities (13). Therefore, we sought to uncover relationships between the two identified clusters and eating behaviours assessed using the BEBQ. Spearman correlations were calculated between the relative abundances of Clusters I and II, and the five eating behaviours, and then visualized with a heatmap. Cluster I relative abundance was significantly correlated with food responsiveness (R = −0.23, P = 0.03; Figure 3c), and both clusters were significantly correlated with enjoyment of food (Cluster I: R = 0.22, P =0.04; Cluster II: R = −0.23, P = 0.03; Figure 3a, Supplementary Figure 5). Food responsiveness is significantly higher in breastfed infants compared to formula-fed infants (P = 0.039, Figure 3b), aligning with literature (6). These findings lead us to postulate that Cluster I relative abundance, which is regulated by diet, is inversely associated with food responsiveness.

**FIG. 3.**
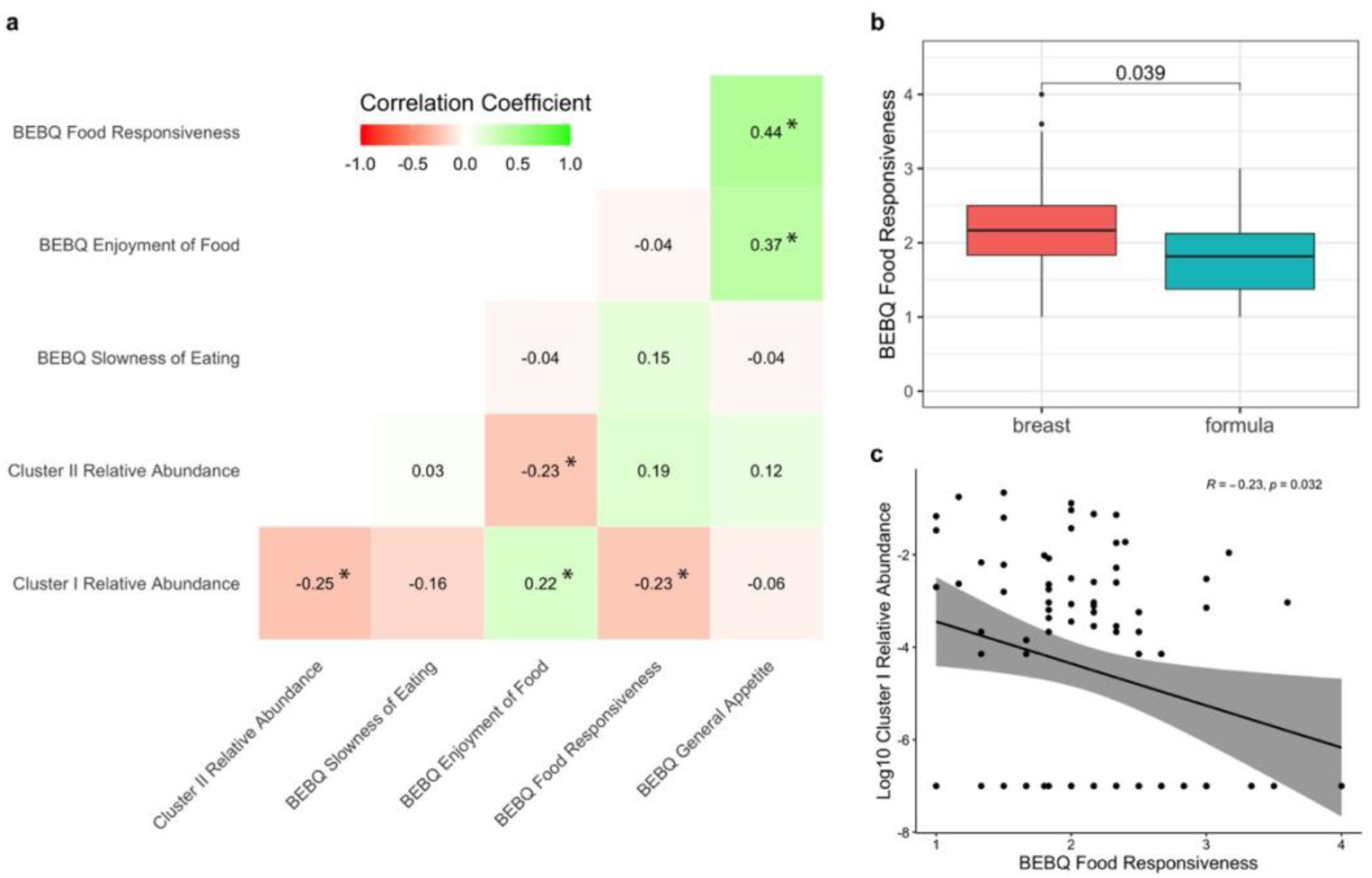
Diet is a driver of the relationships between food responsiveness and relative cluster abundance. (A) Heatmap of correlation matrix between cluster abundance and BEBQ behaviour metrics coloured by Spearman correlation coefficient. Significance (p < 0.05) is marked with a star (*). (B, C) Spearman correlation coefficient and significance for cluster I and food responsiveness for breastfed and formula-fed infants combined and separately.

### Fermentation of succinate to butanoate is significantly negatively correlated with food responsiveness

Finally, we sought to identify metabolic pathways that link Cluster I with food responsiveness. Using ALDeX2 to analyse metabolic pathways inferred by PICRUSt2, six pathways were found to be significantly differentially expressed between breastfed and formula-fed infants (Supplementary Table 2, Figure 4a). Two involve fermentation to short-chain fatty acids (SCFAs): acetyl-CoA fermentation to butanoate II and succinate fermentation to butanoate. SCFAs are important in the context of gut microbiota and eating behaviours because they regulate food intake and are associated with alterations in body weight (40). Both are significantly more abundant in formula-fed infants than breastfed infants (P < 0.001 for all three; Supplementary Figure 6a, Figure 4b).

**FIG. 4.**
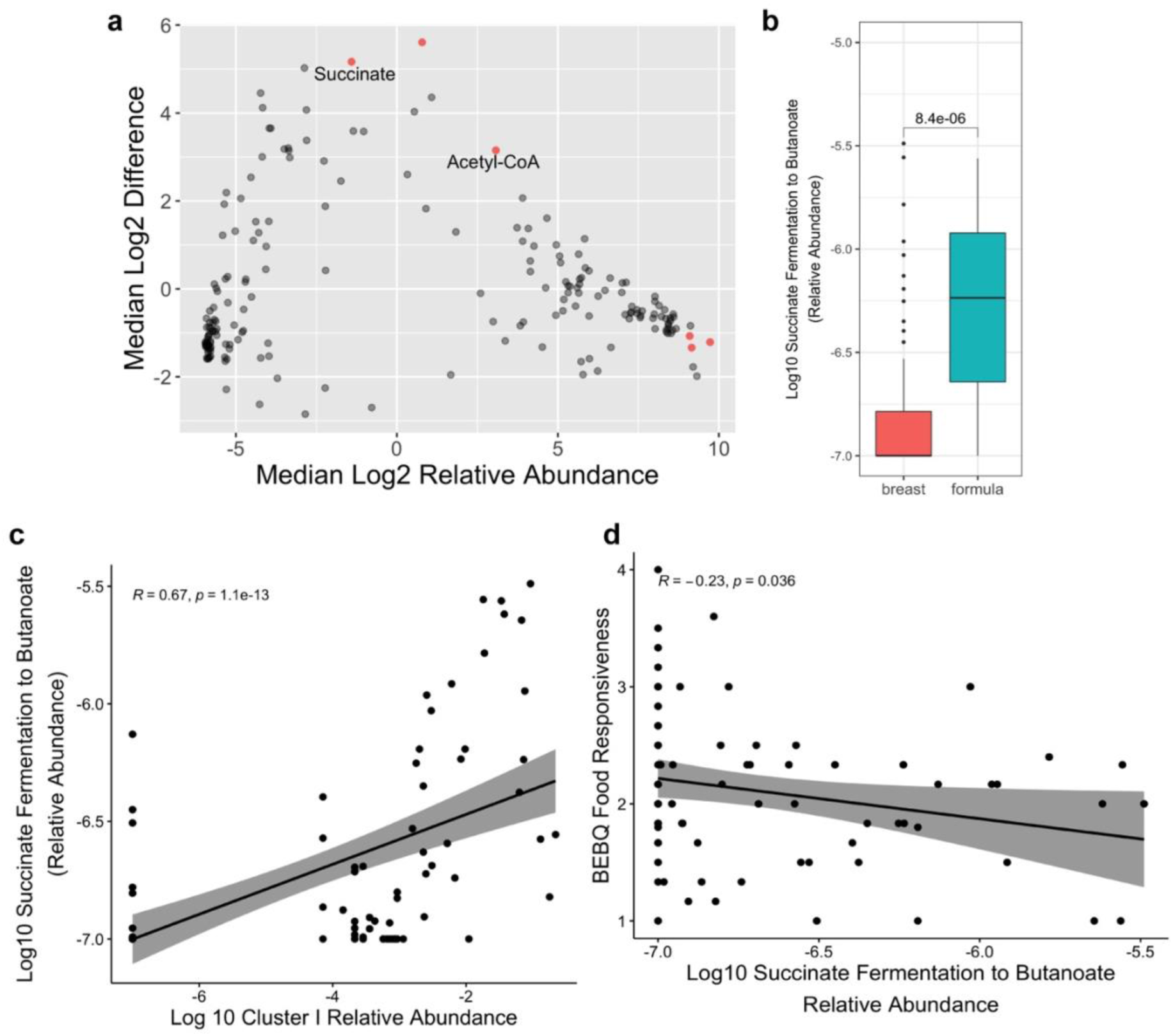
Fermentation of succinate to butanoate is significantly negatively correlated with food responsiveness. (A) Metabolic pathways within breastfed and formula-fed infants inferred using PICRUSt2. Differentially abundant pathways (p < 0.05 for Benjamini-Hochberg corrected P values for Welch’s t-test and Wilcoxon test) are coloured in red. Two differentially abundant pathways relevant to SCFA synthesis (acetyl-CoA fermentation to butanoate II, and succinate fermentation to butanoate) are labelled. (B) Succinate fermentation to butanoate is significantly more highly expressed in formula-fed infants according to the Mann-Whitney U test. (C, D) Succinate fermentation to butanoate is significantly positively correlated with cluster I abundance and negatively correlated with food responsiveness based on Spearman correlation coefficient and p-values.

While the relative abundances of both are significantly positively correlated with Cluster I relative abundance (R = 0.54 and 0.67, respectively and P < 0.001 for both), none are significantly correlated with Cluster II relative abundance (R = −0.19 and −0.08 and P = 0.07 and 0.41, respectively; Supplementary Figure 6b, Figure 4c). These data align with how the phylum Firmicutes is commonly responsible for SCFA production (41). Only one, succinate fermentation to butanoate, is significantly negatively correlated with food responsiveness (R = −0.23, P = 0.04; Figure 4d). Due to the role of SCFAs in appetite suppression, this result is expected (42). The production of SCFAs results in appetite suppression through increasing anorectic gut hormones like glucagon-like peptide 1 (GLP-1) (43). GLP-1 induces satiety and reduces weight gain by increasing insulin secretion following food intake (44). As such, our results seem to suggest that higher abundances of Cluster I in formula-fed infants may decrease the food responsiveness of these infants through SCFA production.

One study suggests that breastfeeding protects against childhood obesity (45), which contradicts our suggested model. An explanation for this contradiction is that high abundances of SCFA-producers, such as Firmicutes, in the stool of formula-fed infants do not always indicate proper SCFA absorption by the infant. SCFA and metabolites may be excreted instead of absorbed, reducing satiety and increasing the risk of obesity (43). Furthermore, prior studies are discrepant with regard to whether breastfed or formula-fed infants show greater SCFA production and absorption, and report that SCFA distributions in infants vary by infant age (43). There is also no literature regarding how SCFA correlates with infant weight and whether breastfeeding protects against childhood obesity remains contested. Therefore, our proposed model, rather than being a dogma, should be treated as a call to further research regarding the relationship between the infant’s diet, gut microbiome, and eating behaviours.

## CONCLUSION

Our study investigated how the infant’s diet impacts their gut microbiota and eating behaviours during the first 4 months of life. We initially hypothesized that formula-feeding would be associated with a higher abundance of the phylum Firmicutes and exhibition of obesity-prone eating behaviours. While we did find that formula-fed infants hosted greater levels of a cluster rich in Firmicutes, these microbes were associated with lower levels of food responsiveness, which would theoretically correspond to a lower obesity risk. Furthermore, we identified the production of SCFAs by the Firmicutes-rich cluster as a mechanism for decreasing food responsiveness. Our model postulates that formula impacts eating behaviours by altering

SCFA-producers in the gut microbiome. How these microbes change over time, prime the gut for future microbial colonization, and affect longer-term eating behaviors and growth trajectories remain to be seen. However, these results provide a new understanding of the psychophysiological impacts of gut microbial communities in the first 4 months of life, and calls for additional research to be done to better understand how infant diet impacts the development of adult microbial communities and subseqnet growth and development of eating behaviors. Greater understanding of these factors can potentially inform strategies for childhood obesity prevention.

## Supporting information

Supplemental R Markdown

## ACKNOWLEDGMENTS

We thank the MICB 447 instructors Dr. Dave Oliver, Dr. Stephan Koenig, Emily Adamcyzk, and Mihai Cirstea at the University of British Columbia for their guidance. We would also like to acknowledge the UC San Diego Center for Microbiome Innovation for their collaboration in sequencing the samples. This work was supported by American Heart Association [grant number 13EIA14660045] and the Eunice Kennedy Shriver National Institute of Child Health and Human Development [R01HD084163].

