## Supplemental R Markdown for "Diet Potentially Drives the Differentiation of Eating Behaviours via Alterations to the Gut Microbiome in Infants"

### Supplementary: Code for Analyses and Figures

Cathy Yan

23/03/2021

```
knitr::opts_chunk$set(echo = TRUE)
set.seed(4321)
```

#### Set-up

Loading packages:

```
library(phyloseq)
library(DESeq2)
library(tidyverse)
library(data.table)
library(ggpubr)
library(caret)
library(randomForest)
library(ROCR)
library(compositions)
library(Hmisc)
library(ALDEx2)
library(corrplot)
library(ape)
library(btools)
library(vegan)
```

Creating a phyloseq object:

```
biom_file <- import_biom("new_tax_table.biom") # taxonomic composition for each sample
```

```
## Warning in strsplit(conditionMessage(e), "\n"): input string 1 is invalid in
## this locale
```

```
metadata <- import_qiime_sample_data("infant_metadata2.txt") # metadata
tree <- read_tree_greengenes("tree.nwk")
tree <- multi2di(tree)

physeq <- merge_phyloseq(biom_file, metadata, tree)

#Add taxonomic rank names
colnames(tax_table(physeq)) <- c("Kingdom", "Phylum", "Class", "Order", "Family", "Genus", "Species")

# Load in metadata data frame
meta <- read_tsv("infant_metadata2.txt")
```

```
##
## -- Column specification --------------------------------------------------------
## cols(
##   .default = col_character(),
##   anonymized_name = col_double(),
##   host_subject_id = col_double(),
##   timepoint = col_double(),
##   waz_avg = col_double(),
##   wlz_avg = col_double()
## )
## i Use `spec()` for the full column specifications.
```

```
#Rarefy at 14000 reads
physeq_rar <- rarefy_even_depth(physeq, sample.size = 14000, rngseed = TRUE) %>%
  subset_samples(feed %in% c("breast", "formula") & age_category %in% c("0.5 months", "2 months", "4 mon
```

```
## `set.seed(TRUE)` was used to initialize repeatable random subsampling.

## Please record this for your records so others can reproduce.

## Try `set.seed(TRUE); .Random.seed` for the full vector

## ...

## 17 samples removedbecause they contained fewer reads than `sample.size`.

## Up to first five removed samples are:

## 10918.38693.R1.fastq.gz10918.38697.R1.fastq.gz10918.38703.R1.fastq.gz10918.38705.R1.fastq.gz10918.387

## ...

## 140OTUs were removed because they are no longer
## present in any sample after random subsampling

## ...
```

```
physeq_rar_RA <- transform_sample_counts(physeq_rar, function(x) x/sum(x)) # relative abundance
```

#### Participant Characteristics

##### Creating Data Frames for Later Use

```
#Make genus abundance table
otu_table <- as.data.frame(otu_table(physeq_rar_RA)) %>%
  setDT(keep.rownames="OTUID") #sample names as columns, otu ID's as rows
tax_table <- as.data.frame(tax_table(physeq_rar_RA)) %>%
  setDT(keep.rownames="OTUID") %>%
```

```
  dplyr::select(OTUID, Genus) %>%
  filter(is.na(Genus) == F, Genus != "g__")
tax_table$Genus <- gsub("\\[|\\]", "", tax_table$Genus) #removing the square brackets around some names

#Merge with taxonomies
genus_ab_table <- merge(tax_table, otu_table) %>%
  dplyr::select(-OTUID) %>%
  group_by(Genus) %>%
  dplyr::summarize(across(everything(), list(sum))) %>%
  column_to_rownames(var="Genus")
```

```
## `summarise()` ungrouping output (override with `.groups` argument)
```

```
#Merge with metadata
transposed_rel_ab <- t(genus_ab_table) %>%
  as.data.frame() %>%
  setDT(keep.rownames="#SampleID")
transposed_rel_ab$`#SampleID` <- gsub("_1", "", transposed_rel_ab$`#SampleID`)

feed_meta <- meta %>%
  dplyr::select("#SampleID", feed) #starting with feed

feed_meta_abundance <- inner_join(transposed_rel_ab, feed_meta) %>%
  dplyr::select(-`#SampleID`) %>%
  filter(feed %in% c("breast", "formula"))
```

```
## Joining, by = "#SampleID"
```

```
feed_meta_abundance$feed <- as.factor(feed_meta_abundance$feed)

filtered_meta <- meta %>%
  filter(`#SampleID` %in% transposed_rel_ab$`#SampleID`)
```

##### Getting Information on Eating Behaviours

```
# Number of infants
length(unique(filtered_meta$anonymized_name))
```

```
## [1] 58
```

```
# Breastfed infants
nrow(filter(filtered_meta, feed == "breast"))
```

```
## [1] 82
```

```
filtered_meta$bebq_enjoyment_food <- as.numeric(filtered_meta$bebq_enjoyment_food)
```

```
## Warning: NAs introduced by coercion
```

```
filtered_meta$bebq_food_responsive <- as.numeric(filtered_meta$bebq_food_responsive)
```

```
## Warning: NAs introduced by coercion
```

```
filtered_meta$bebq_gen_appetite <- as.numeric(filtered_meta$bebq_gen_appetite)
```

```
## Warning: NAs introduced by coercion
```

```
filtered_meta$bebq_slowness_eat <- as.numeric(filtered_meta$bebq_slowness_eat)
```

```
## Warning: NAs introduced by coercion
```

```
# Eating behaviour mean/median
stats <- filtered_meta %>%
  filter(age_category == "4 months") %>% # Options are 0.5 months, 2 months, and 4 months
  dplyr::select(feed, bebq_enjoyment_food, bebq_food_responsive, bebq_gen_appetite, bebq_slowness_eat) %
  as.data.frame() %>%
  gather("metric", "score", -feed)  %>%
  group_by(feed, metric) %>%
  dplyr::summarize(mean = mean(score, na.rm = TRUE),
                   median = median(score, na.rm = TRUE),
                   SD = sd(score, na.rm = TRUE),
                   count = n())
```

```
## `summarise()` regrouping output by 'feed' (override with `.groups` argument)
```

```
stats
```

```
## # A tibble: 8 x 6
## # Groups:   feed [2]
##   feed    metric                mean median    SD count
##   <chr>   <chr>                <dbl>  <dbl> <dbl> <int>
## 1 breast  bebq_enjoyment_food   4.29   4.38 0.570    20
## 2 breast  bebq_food_responsive  1.94   1.83 0.624    20
## 3 breast  bebq_gen_appetite     3.67   4    0.907    20
## 4 breast  bebq_slowness_eat     2.53   2.62 0.664    20
## 5 formula bebq_enjoyment_food   4.58   4.62 0.342     6
## 6 formula bebq_food_responsive  1.75   1.83 0.673     6
## 7 formula bebq_gen_appetite     3.83   4    0.753     6
## 8 formula bebq_slowness_eat     2      2.12 0.524     6
```

#### Alpha and Beta Diversity Analyses

##### Beta Diversity

```
# Calculating distances
wunifrac_dist <- phyloseq::distance(physeq_rar_RA, method="wunifrac")
unifrac_dist <- phyloseq::distance(physeq_rar_RA, method="unifrac")
```

```r
jaccard_dist <- phyloseq::distance(physeq_rar_RA, method="jaccard")
bray_dist <- phyloseq::distance(physeq_rar_RA, method="bray")

ordw <- ordinate(physeq_rar_RA, method = "PCoA",
                 distance = "wunifrac")
ordj <- ordinate(physeq_rar_RA, method = "PCoA",
                 distance = "jaccard")
ordu <- ordinate(physeq_rar_RA, method = "PCoA",
                 distance = "unifrac")
ordb <- ordinate(physeq_rar_RA, method = "PCoA",
                 distance = "bray")

# Assessing confounders
adonis(wunifrac_dist ~ sample_data(physeq_rar_RA)$abx_any_source)
```

```
##
## Call:
## adonis(formula = wunifrac_dist ~ sample_data(physeq_rar_RA)$abx_any_source)
##
## Permutation: free
## Number of permutations: 999
##
## Terms added sequentially (first to last)
##
##                                          Df SumsOfSqs  MeanSqs F.Model      R2
## sample_data(physeq_rar_RA)$abx_any_source  1    0.1220 0.122013  1.4102 0.01478
## Residuals                                94    8.1333 0.086525         0.98522
## Total                                    95    8.2554                  1.00000
##                                          Pr(>F)
## sample_data(physeq_rar_RA)$abx_any_source  0.217
## Residuals
## Total
```

```r
adonis(wunifrac_dist ~ sample_data(physeq_rar_RA)$baby_delivery)
```

```
##
## Call:
## adonis(formula = wunifrac_dist ~ sample_data(physeq_rar_RA)$baby_delivery)
##
## Permutation: free
## Number of permutations: 999
##
## Terms added sequentially (first to last)
##
##                                         Df SumsOfSqs  MeanSqs F.Model      R2
## sample_data(physeq_rar_RA)$baby_delivery  2    0.2859 0.142973  1.6684 0.03464
## Residuals                               93    7.9694 0.085693         0.96536
## Total                                   95    8.2554                  1.00000
##                                         Pr(>F)
## sample_data(physeq_rar_RA)$baby_delivery  0.122
## Residuals
## Total
```

```r
adonis(wunifrac_dist ~ sample_data(physeq_rar_RA)$pbx_any_source)
```

```
##
## Call:
## adonis(formula = wunifrac_dist ~ sample_data(physeq_rar_RA)$pbx_any_source)
##
## Permutation: free
## Number of permutations: 999
##
## Terms added sequentially (first to last)
##
##                                           Df SumsOfSqs  MeanSqs F.Model      R2
## sample_data(physeq_rar_RA)$pbx_any_source  1    0.0483 0.048269 0.55285 0.00585
## Residuals                                 94    8.2071 0.087309         0.99415
## Total                                     95    8.2554                  1.00000
##                                           Pr(>F)
## sample_data(physeq_rar_RA)$pbx_any_source   0.67
## Residuals
## Total
```

```r
# Supplementary Figure 1
plot_ordination(physeq_rar_RA,
                ordb,
                type = "sample",
                color = "feed",
                title = "") +
  stat_ellipse(type = "norm", size = 1) +
  theme(axis.text = element_text(size = 12),
        axis.title = element_text(size = 16),
        legend.text = element_text(size = 12),
        legend.title = element_text(size = 16)) +
  labs(color = "Diet")
```

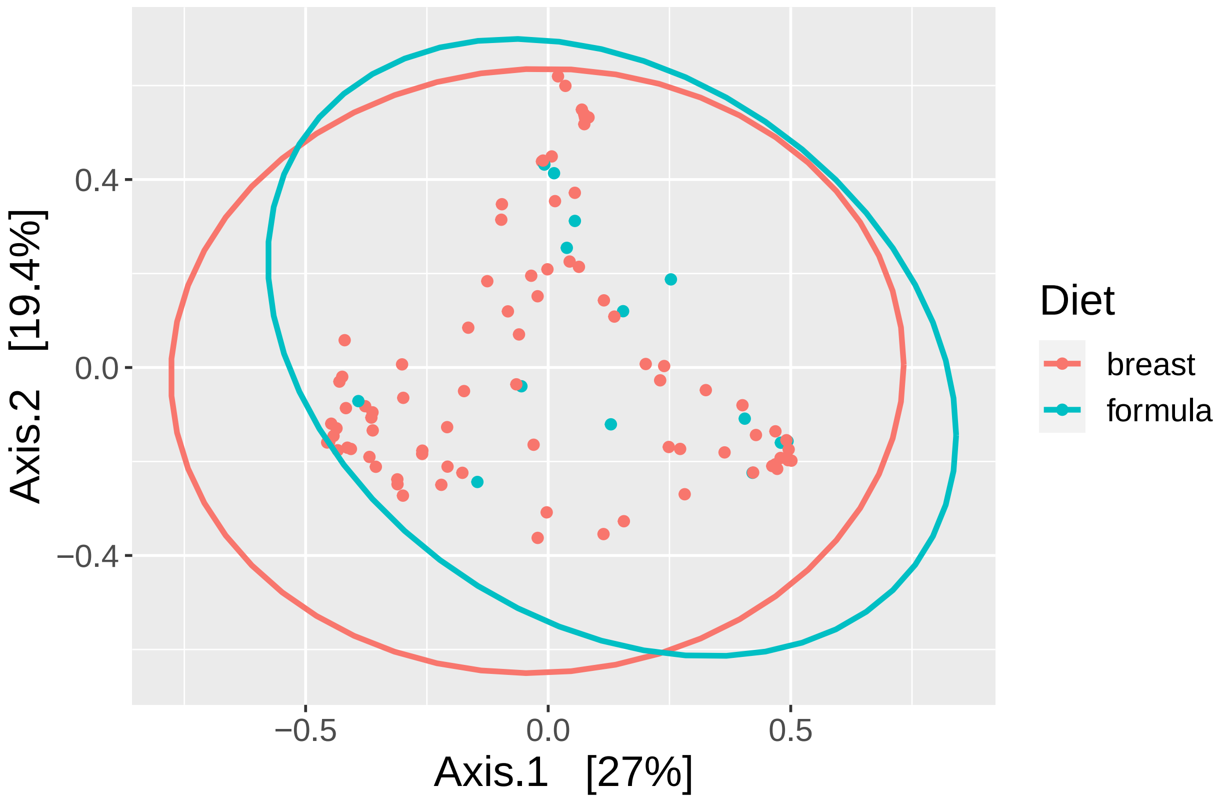


```r
# ggsave("Figures/brayplot.png", width = 8, height = 4, dpi = 600)

plot_ordination(physeq_rar_RA,
                ordj,
                type = "sample",
                color = "feed",
                title = "") +
  stat_ellipse(type = "norm", size = 1) +
  theme(axis.text = element_text(size = 12),
        axis.title = element_text(size = 16),
        legend.text = element_text(size = 12),
        legend.title = element_text(size = 16)) +
  labs(color = "Diet")
```

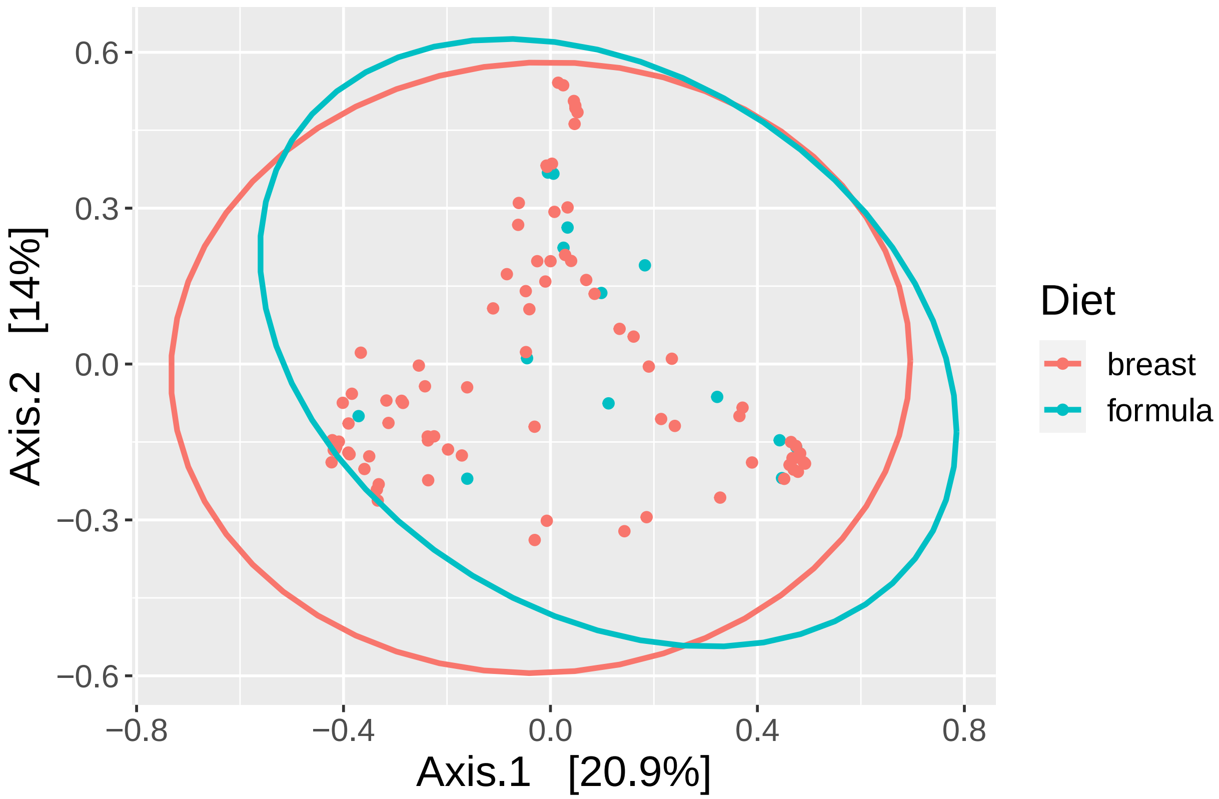


```r
# ggsave("Figures/jaccardplot.png", width = 8, height = 4, dpi = 600)

plot_ordination(physeq_rar_RA,
                ordu,
                type = "sample",
                color = "feed",
                title = "") +
  stat_ellipse(type = "norm", size = 1) +
  theme(axis.text = element_text(size = 12),
        axis.title = element_text(size = 16),
        legend.text = element_text(size = 12),
        legend.title = element_text(size = 16)) +
  labs(color = "Diet")
```

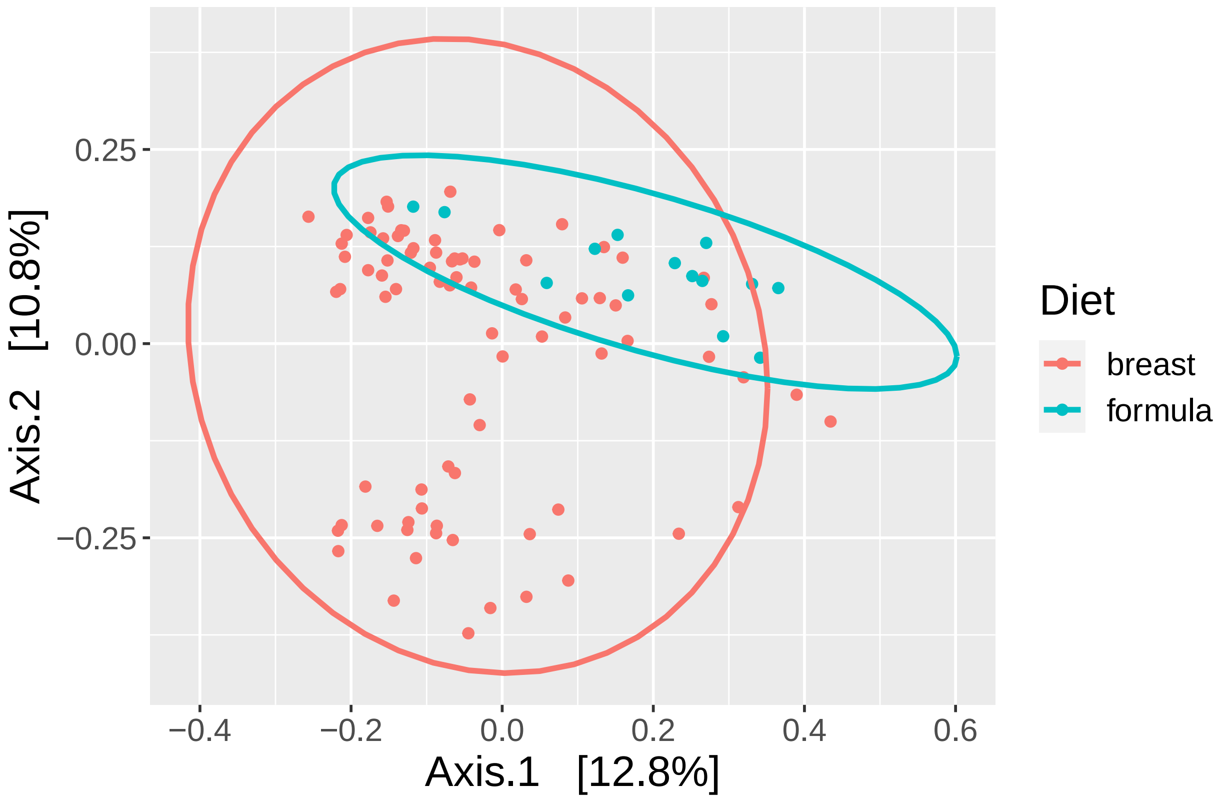


```
# ggsave("Figures/unifracplot.png", width = 8, height = 4, dpi = 600)

# Figure 1a
plot_ordination(physeq_rar_RA,
                ordw, #we can use the same distance calculations as above
                type = "sample",
                color = "feed",
                title = "") +
  stat_ellipse(type = "norm", size = 1) +
  theme(axis.text = element_text(size = 12),
        axis.title = element_text(size = 16),
        legend.text = element_text(size = 12),
        legend.title = element_text(size = 16)) +
  labs(color = "Diet")
```

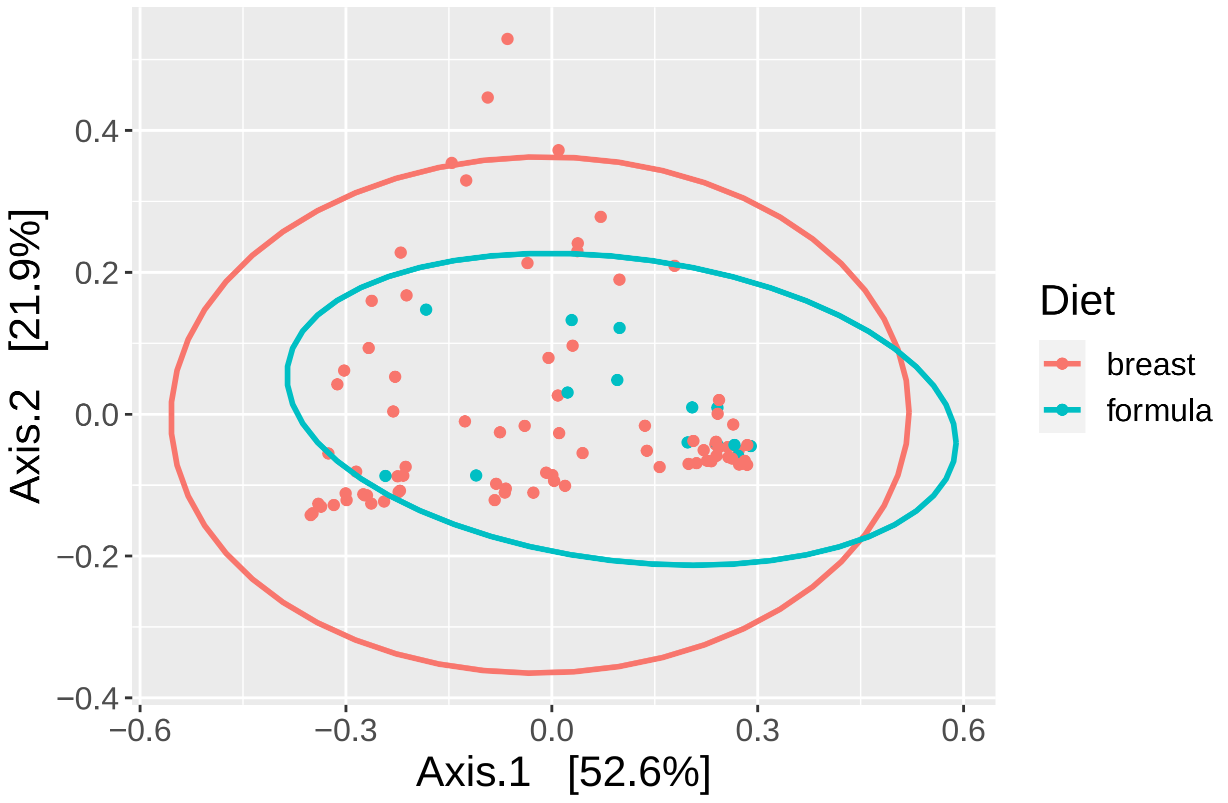


```r
# ggsave("UpdatedFigs/wunifracplot.png", width = 8, height = 4, dpi = 600)

# PERMANOVA
adonis(wunifrac_dist ~ sample_data(physeq_rar_RA)$feed)
```

```
##
## Call:
## adonis(formula = wunifrac_dist ~ sample_data(physeq_rar_RA)$feed)
##
## Permutation: free
## Number of permutations: 999
##
## Terms added sequentially (first to last)
##
##                                 Df SumsOfSqs  MeanSqs F.Model      R2 Pr(>F)
## sample_data(physeq_rar_RA)$feed  1    0.3117 0.311661   3.688 0.03775   0.02 *
## Residuals                       94    7.9437 0.084507         0.96225
## Total                           95    8.2554                  1.00000
## ---
## Signif. codes:  0 '***' 0.001 '**' 0.01 '*' 0.05 '.' 0.1 ' ' 1
```

**Alpha Diversity**

```
# Alpha diversity supplementary plot
alpha <- plot_richness(physeq_rar, "feed") #we want box plots instead

feed_comp <- list(c("breast", "formula"))

a_dat <- alpha$data
a_plot <- ggplot(a_dat, aes(x = feed, y = value)) +
  geom_boxplot() +
  facet_wrap(~variable, scales = "free") +
  xlab("Diet") +
  ylab("Value") +
  stat_compare_means(comparisons = feed_comp, method = "wilcox.test",
                     label = "p.format", label.y.npc = 0.75, label.x = 1.3, size = 3)
a_plot
```

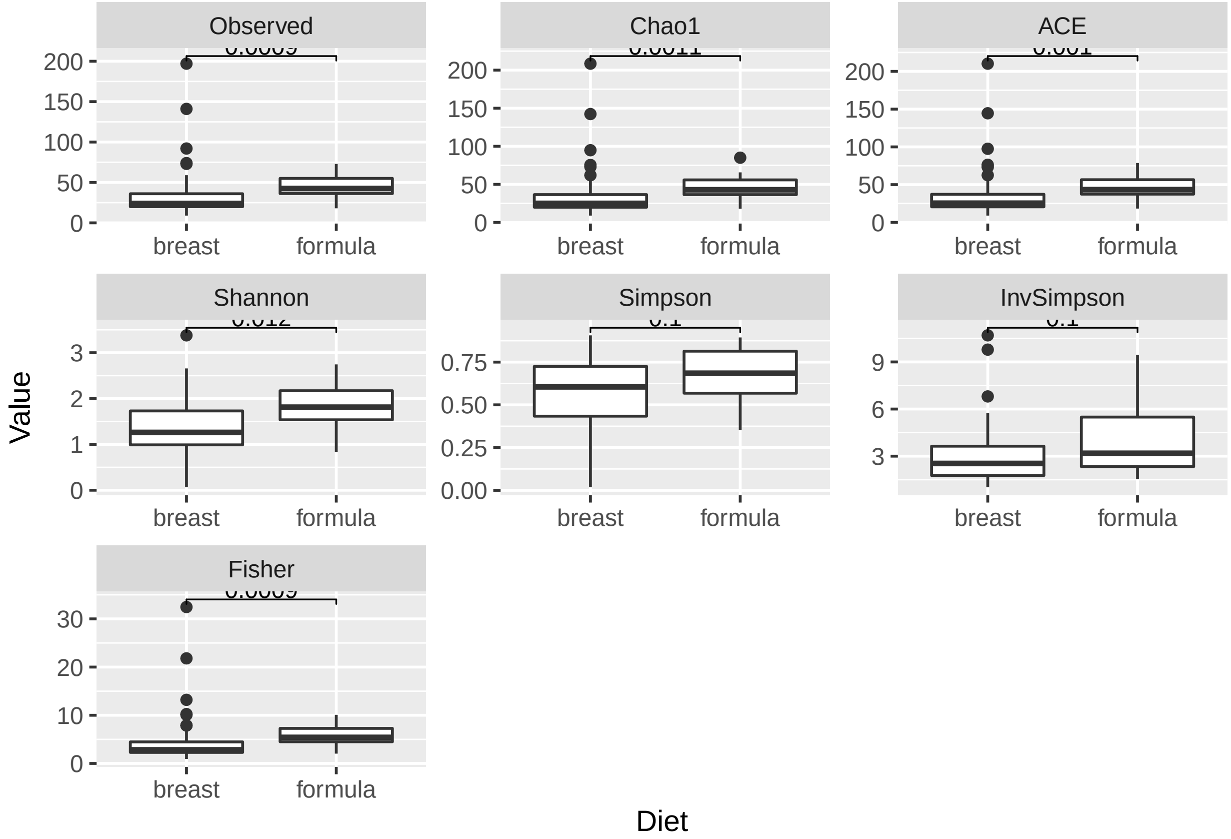


```
# ggsave("Figures/Supp3.png", plot = a_plot, width = 8, height = 12, dpi = 300)

# Figure 1b
faith <- estimate_pd(physeq_rar) %>%
  setDT(keep.rownames="#SampleID") %>%
  inner_join(meta)
```

```
## Calculating Faiths PD-index...

## Joining, by = "#SampleID"
```

```r
faith_plot <- ggplot(faith, aes(x = as.factor(feed), y = PD)) +
  geom_boxplot() +
  xlab("Diet") +
  ylab("Faith's Phylogenetic Diversity") +
  ylim(0, 20) +
  theme(axis.text = element_text(size = 12),
        axis.title = element_text(size = 16)) +
  stat_compare_means(comparisons = feed_comp, method = "wilcox.test",
                     label = "p.format", label.y = 19, label.x = 1.3, size = 6)
faith_plot
```

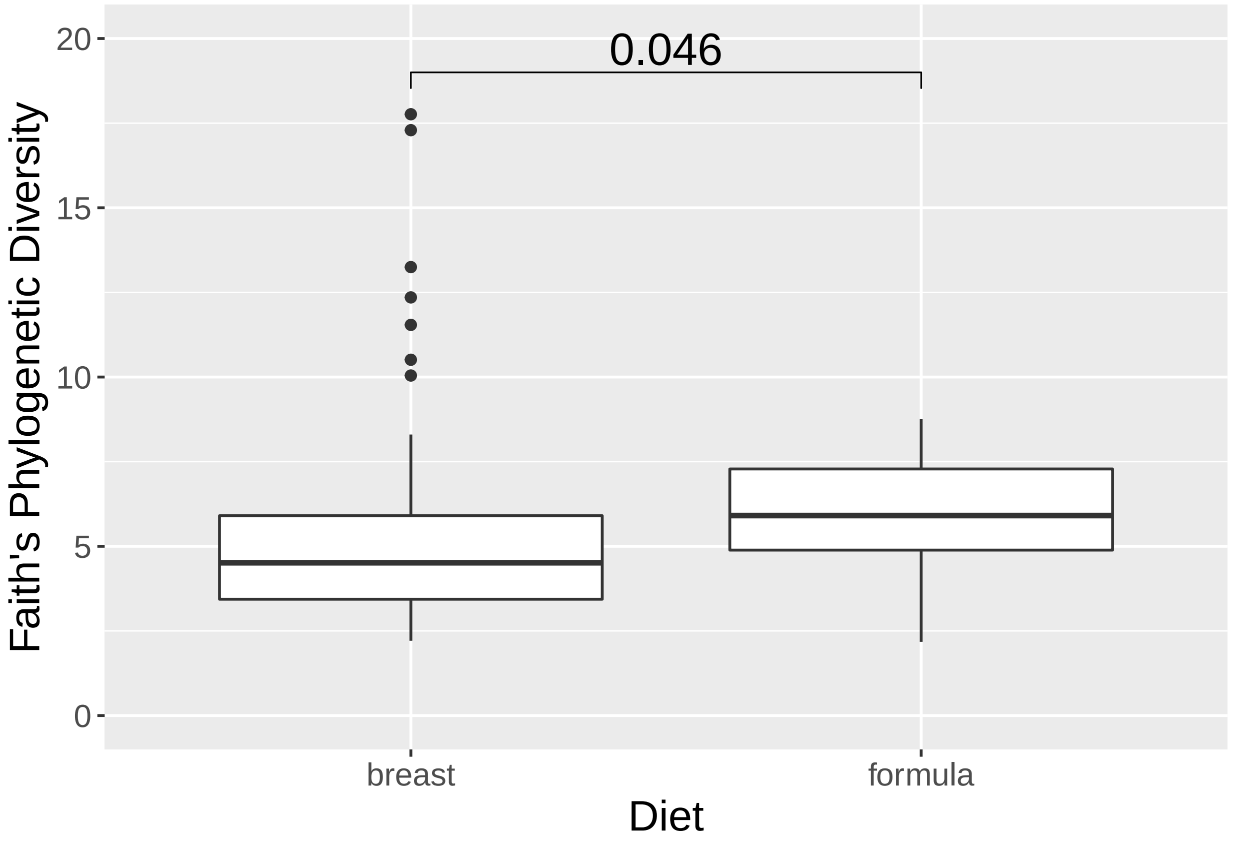


```r
# ggsave("UpdatedFigs/faithplot.jpg", plot = faith_plot, width = 3, height = 5, dpi = 600)
```

##### Random Forest Classifier

```r
set.seed(5619)
#splitting into training and test
rows <- feed_meta_abundance %>%
  dplyr::select(feed) %>%
  unlist() %>%
  createDataPartition(p=0.6, list=FALSE)

feed_train <- feed_meta_abundance %>% slice(rows)
```

```r
feed_test <- feed_meta_abundance %>% slice(-rows)

#set up cross-validation
trControl <- trainControl(method='cv', number=10, search='grid')
tuneGrid <- expand.grid(.mtry = 10)

#final model - tuning steps omitted for brevity
fit_rf <- train(feed~.,
                data=feed_train,
                method = "rf",
                metric = "Accuracy",
                tuneGrid = tuneGrid,
                trControl = trControl,
                importance = TRUE,
                nodesize = 5,
                ntree = 600,
                maxnodes = 15)
```

```
## Warning in nominalTrainWorkflow(x = x, y = y, wts = weights, info = trainInfo, :
## There were missing values in resampled performance measures.
```

```r
predictions <- predict(fit_rf, feed_test)
confusionMatrix(predictions, feed_test$feed)
```

```
## Confusion Matrix and Statistics
##
##           Reference
## Prediction breast formula
##    breast      32       5
##    formula      0       0
##
##                Accuracy : 0.8649
##                  95% CI : (0.7123, 0.9546)
##     No Information Rate : 0.8649
##     P-Value [Acc > NIR] : 0.61627
##
##                   Kappa : 0
##
##  Mcnemar's Test P-Value : 0.07364
##
##             Sensitivity : 1.0000
##             Specificity : 0.0000
##          Pos Pred Value : 0.8649
##          Neg Pred Value :    NaN
##              Prevalence : 0.8649
##          Detection Rate : 0.8649
##    Detection Prevalence : 1.0000
##       Balanced Accuracy : 0.5000
##
##        'Positive' Class : breast
##
```

```r
varImp(fit_rf)
```

```
## rf variable importance
##
##   only 20 most important variables shown (out of 162)
##
##                                  Importance
## g__Blautia                           100.00
## g__Eubacterium                        80.09
## g__Dorea                              59.12
## g__Akkermansia                        56.58
## g__Eggerthella                        53.22
## g__Haemophilus                        49.08
## g__Coprococcus                        48.04
## g__Acinetobacter                      46.38
## g__Proteus                            44.50
## g__Ruminococcus                       43.07
## g__Pseudoramibacter_Eubacterium       40.00
## g__Megasphaera                        39.67
## g__Coprobacillus                      39.00
## g__Porphyromonas                      37.81
## g__Clostridium                        34.81
## g__Anaerofustis                       34.12
## g__Anaerotruncus                      27.47
## g__Oscillospira                       26.27
## g__Anaerococcus                       26.12
## g__Collinsella                        25.69
```

```r
#ROC curve
prediction_for_roc_curve <- predict(fit_rf,feed_test,type="prob")

# Specify the different classes
classes <- levels(feed_test$feed)

true_values_b <- ifelse(feed_test$feed=="breast",1,0)
true_values_f <- ifelse(feed_test$feed=="formula",1,0)

pred_b <- prediction(prediction_for_roc_curve[,1],true_values_b)
pred_f <- prediction(prediction_for_roc_curve[,2],true_values_f)

perf_b <- performance(pred_b, "sens", "spec")
auc.perf_b <- performance(pred_b, measure = "auc")
auc.perf_b@y.values
```

```
## [[1]]
## [1] 0.959375
```

```r
perf_b <- data.frame(
  "Specificity" = perf_b@x.values[[1]],
  "Sensitivity" = perf_b@y.values[[1]],
  "Diet" = rep("breast (AUC: 0.96)", 30)
)
```

```r
perf_f <- performance(pred_f, "sens", "spec")
auc.perf_f <- performance(pred_f, measure = "auc")
auc.perf_f@y.values
```

```
## [[1]]
## [1] 0.959375
```

```r
perf_f <- data.frame(
  "Specificity" = perf_f@x.values[[1]],
  "Sensitivity" = perf_f@y.values[[1]],
  "Diet" = rep("formula (AUC: 0.96)", 30)
)

perf <- rbind(perf_b, perf_f)

rfc_plot <- ggplot(perf, aes(x = Specificity, y = Sensitivity, colour = Diet)) +
  geom_path(size = 1) +
  theme(axis.text = element_text(size = 12),
        axis.title = element_text(size = 16),
        legend.text = element_text(size = 12),
        legend.title = element_text(size = 16))
rfc_plot
```

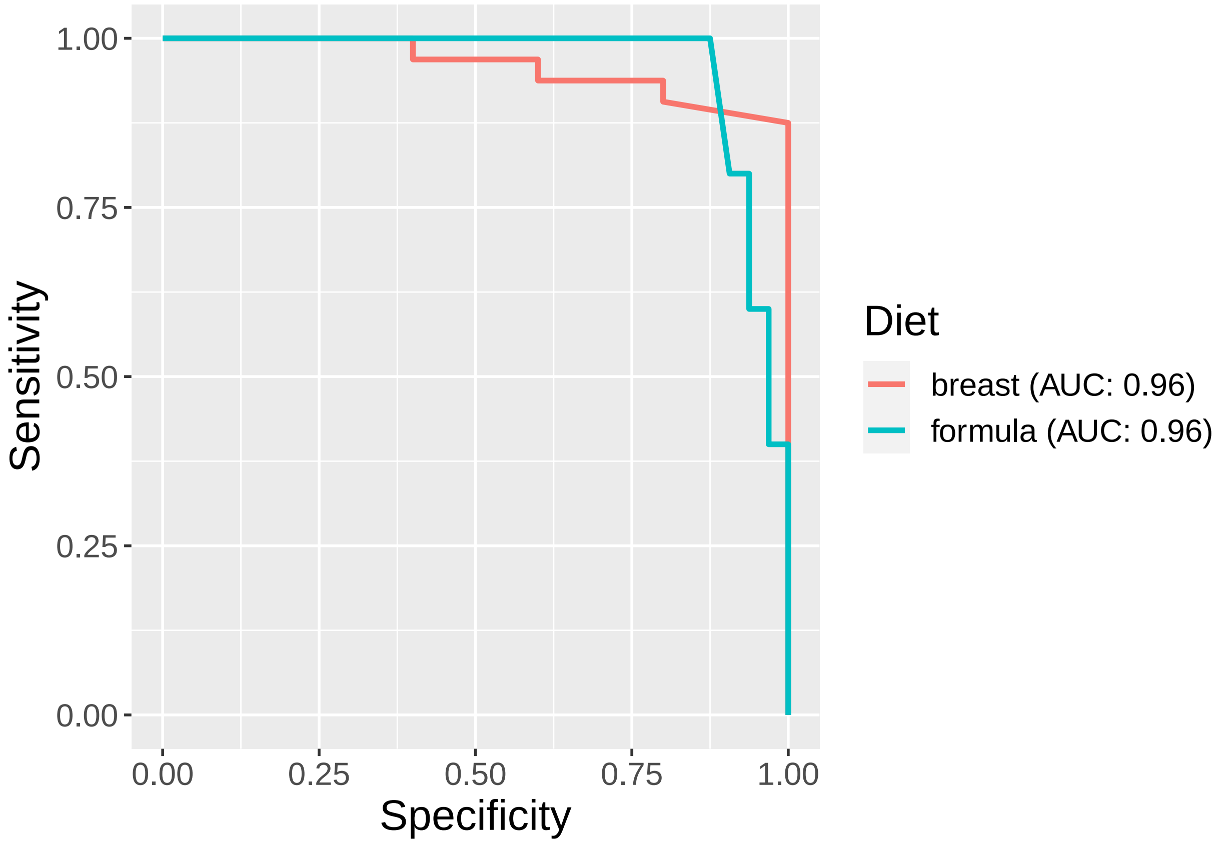

```
# ggsave("randomforestplot.png", plot = rfc_plot, width = 6, height = 4, dpi = 600)
```

#### Co-abundant Clusters

##### Co-abundance Heatmap

```
genus <- transposed_rel_ab %>%
  dplyr::select(-"#SampleID")

genus2 <- genus[, colSums(genus > 0) >= 19] #selecting genera present in at least 20% of the samples
genus3 <- select_if(genus, genus2)

genus1 <- genus3 + 0.0000001 #adding an arbitrarily low value to be able to take the logarithm
genus_clr <- as.data.frame(clr(genus1))
cormat <- cor(genus_clr, method = "spearman")

reorder_cormat <- function(cormat){
  dd <- as.dist(1-cormat)
  hc <- hclust(dd, method = "ward.D")
  cormat <-cormat[hc$order, hc$order]
}

rcormat <- reorder_cormat(cormat)
melted_cormat <- melt(rcormat)
```

```
## Warning in melt(rcormat): The melt generic in data.table has been passed a
## matrix and will attempt to redirect to the relevant reshape2 method; please note
## that reshape2 is deprecated, and this redirection is now deprecated as well.
## To continue using melt methods from reshape2 while both libraries are attached,
## e.g. melt.list, you can prepend the namespace like reshape2::melt(rcormat). In
## the next version, this warning will become an error.
```

```
hc <- hclust(as.dist(1-cor(genus_clr, method="spearman")), method="ward.D")
cut_hc <- cutree(hc, k = 9)
dendro <- plot(hc, xlab = NA)
```

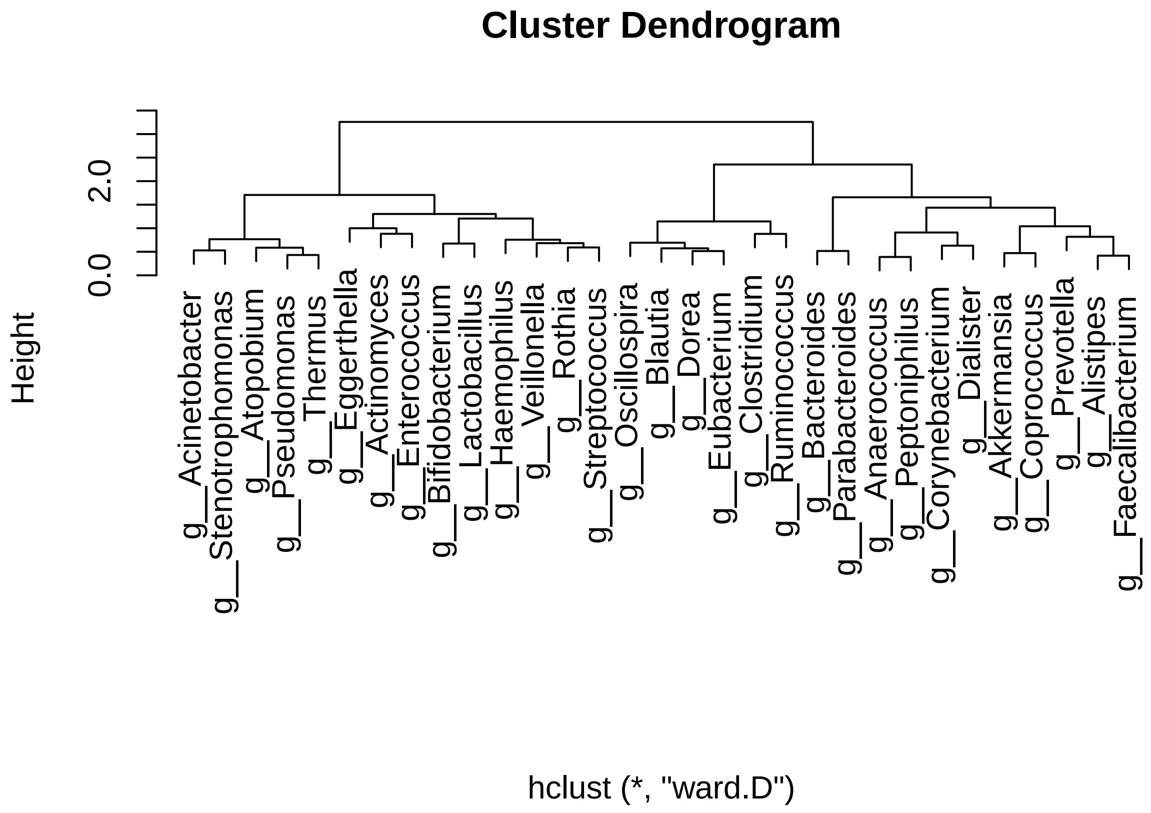


```r
# Figure 2a
ggheatmap <- ggplot(melted_cormat, aes(Var2, Var1, fill = value)) +
  geom_tile(color = "white") +
  scale_fill_gradient2(low = "blue4", high = "darkred", mid = "white",
                       midpoint = 0, limit = c(-0.8,0.8), na.value = "black",
                       space = "Lab", name="Spearman\nCoeff.") +
  theme_minimal() + # minimal theme
  theme(axis.text.x = element_text(angle = 90, vjust = .4, hjust = 1, size = 8),
        axis.text.y = element_text(size = 8)) +
  coord_fixed() +
  ylab("") +
  xlab("")

ggheatmap
```

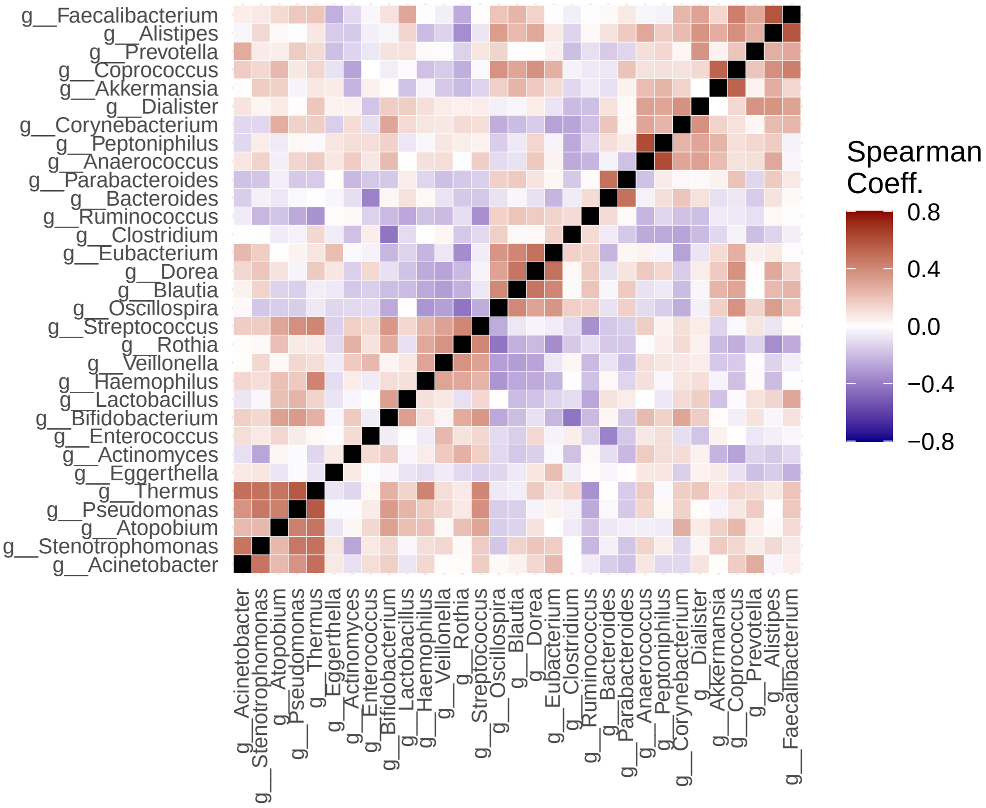


```r
# ggsave("UpdatedFigs/heatmap.png", plot = ggheatmap, width = 6, height = 6, dpi = 600)
```

##### Differentially Abundant Clusters

```r
# Retrieving clusters
cluster_iii <- rcormat[1:5, 1:5]
cluster_ii <- rcormat[9:14, 9:14]
cluster_i <- rcormat[15:18, 15:18]
cluster_iv <- rcormat[23:31, 23:31]

# Calculating relative abundance of each cluster
cluster_1_sum <- transposed_rel_ab %>%
  dplyr::select(row.names(cluster_i)) %>%
  rowSums()
cluster_2_sum <- transposed_rel_ab %>%
  dplyr::select(row.names(cluster_ii)) %>%
  rowSums()
cluster_3_sum <- transposed_rel_ab %>%
  dplyr::select(row.names(cluster_iii)) %>%
  rowSums()
cluster_4_sum <- transposed_rel_ab %>%
  dplyr::select(row.names(cluster_iv)) %>%
  rowSums()
```

```r
# Adding to metadata
filtered_meta <- meta %>%
  filter(`#SampleID` %in% transposed_rel_ab$`#SampleID`)

filtered_meta$clust_i_ab <- cluster_1_sum
filtered_meta$clust_ii_ab <- cluster_2_sum
filtered_meta$clust_iii_ab <- cluster_3_sum
filtered_meta$clust_iv_ab <- cluster_4_sum

filtered_meta <- filter(filtered_meta, feed %in% c("breast", "formula"))

# Plot Comparisons
# Figure 2b
cluster_i_feed <- ggplot(aes(x = as.factor(feed), y = as.numeric(log10(clust_i_ab + 0.0000001))), data =
  geom_boxplot(aes(fill = as.factor(feed))) +
  ylab("Log10 Cluster I Relative Abundance") +
  theme_bw(base_size = 18) +
  xlab(NULL) +
  theme(axis.text.x = element_text(size = 18)) +
  guides(fill = FALSE) +
  ylim(-7, 2) +
  stat_compare_means(comparisons = feed_comp, method = "wilcox.test",
                     label = "p.format", label.y = 0.9, label.x = 1.3, size = 6)
cluster_i_feed
```

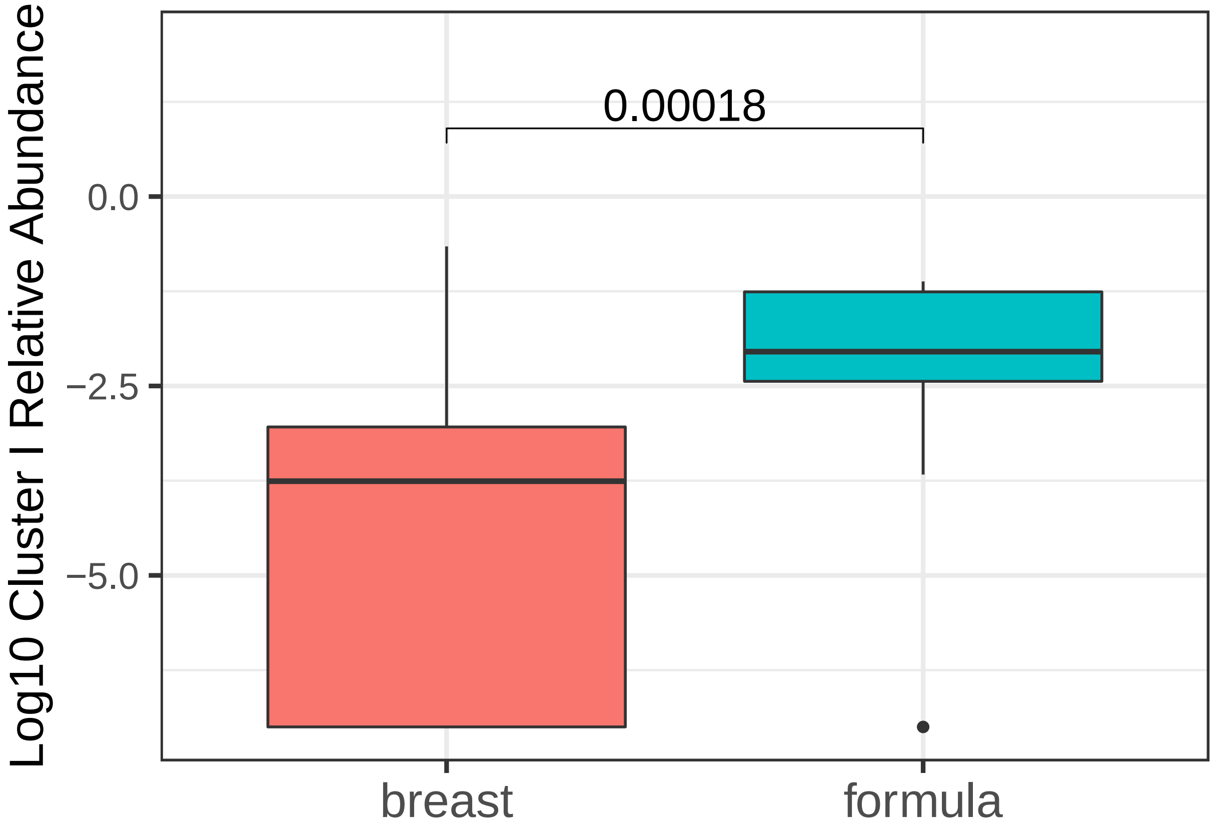

```
# ggsave("UpdatedFigs/clusti.png", plot = cluster_i_feed, width = 4, height = 8, dpi = 600)

# Figure 2c
cluster_ii_feed <- ggplot(aes(x = as.factor(feed), y = log10(clust_ii_ab + 0.0000001)), data = filtered_
  geom_boxplot(aes(fill = as.factor(feed))) +
  ylab("Log 10 Cluster II Relative Abundance") +
  theme_bw(base_size = 18) +
  xlab(NULL) +
  theme(axis.text.x = element_text(size = 18)) +
  guides(fill = FALSE) +
  ylim(-3, 1.5) +
  stat_compare_means(comparisons = feed_comp, method = "wilcox.test",
                     label = "p.format", label.y = 0.9, label.x = 1.3, size = 6)
cluster_ii_feed
```

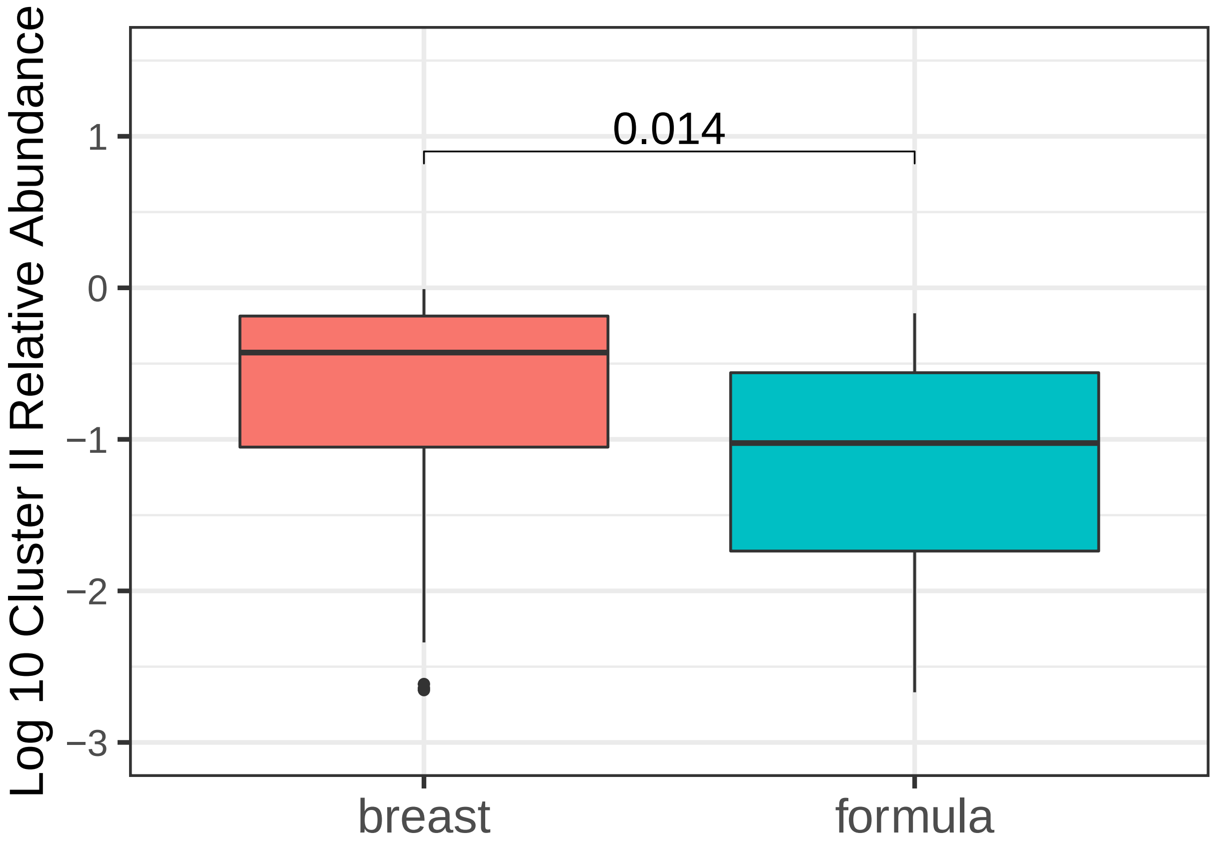


```
# ggsave("UpdatedFigs/clustii.png", plot = cluster_ii_feed, width = 4, height = 8, dpi = 600)

cluster_iii_feed <- ggplot(aes(x = as.factor(feed), y = log10(clust_iii_ab + 0.0000001)), data = filtere
  geom_boxplot(aes(fill = as.factor(feed))) +
  ylab("Log 10 Cluster III Relative Abundance") +
  theme_bw(base_size = 18) +
  xlab(NULL) +
  theme(axis.text.x = element_text(size = 18)) +
  guides(fill = FALSE) +
  ylim(-4, 1) +
```

```r
  stat_compare_means(comparisons = feed_comp, method = "wilcox.test",
                     label = "p.format", label.y = 0.9, label.x = 1.3, size = 6)
cluster_iii_feed
```

```
## Warning: Removed 42 rows containing non-finite values (stat_boxplot).

## Warning: Removed 42 rows containing non-finite values (stat_signif).

## Warning in wilcox.test.default(c(-3.03213791571464, -3.44703644590845, NA, :
## cannot compute exact p-value with ties
```

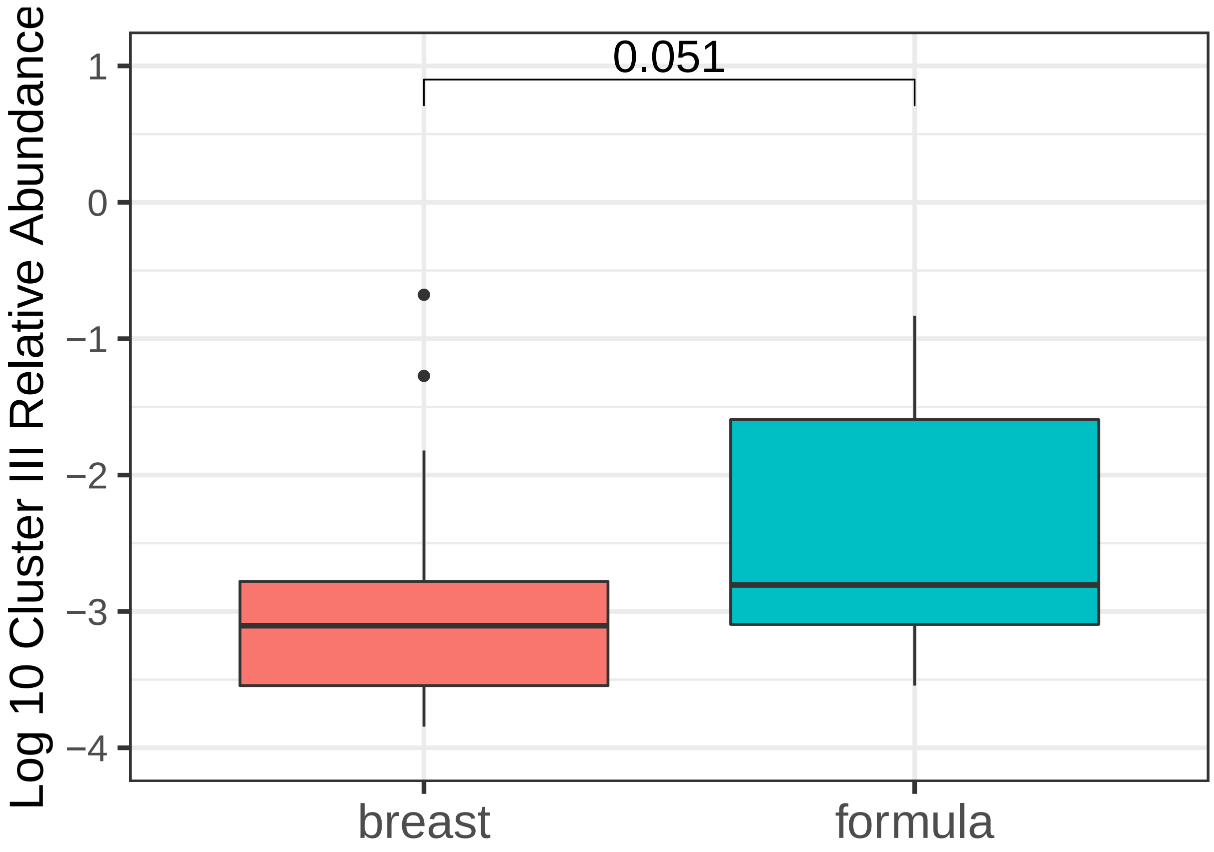


```r
# ggsave("Figures/clustiii.png", plot = cluster_iii_feed, width = 4, height = 8, dpi = 600)

cluster_iv_feed <- ggplot(aes(x = as.factor(feed), y = log10(clust_iv_ab + 0.0000001)), data = filtered_
  geom_boxplot(aes(fill = as.factor(feed))) +
  ylab("Log 10 Cluster IV Relative Abundance") +
  theme_bw(base_size = 18) +
  xlab(NULL) +
  theme(axis.text.x = element_text(size = 18)) +
  guides(fill = FALSE) +
  ylim(-4, 1) +
  stat_compare_means(comparisons = feed_comp, method = "wilcox.test",
                     label = "p.format", label.y = 0.9, label.x = 1.3, size = 6)
cluster_iv_feed
```

```
## Warning: Removed 24 rows containing non-finite values (stat_boxplot).

## Warning: Removed 24 rows containing non-finite values (stat_signif).
```

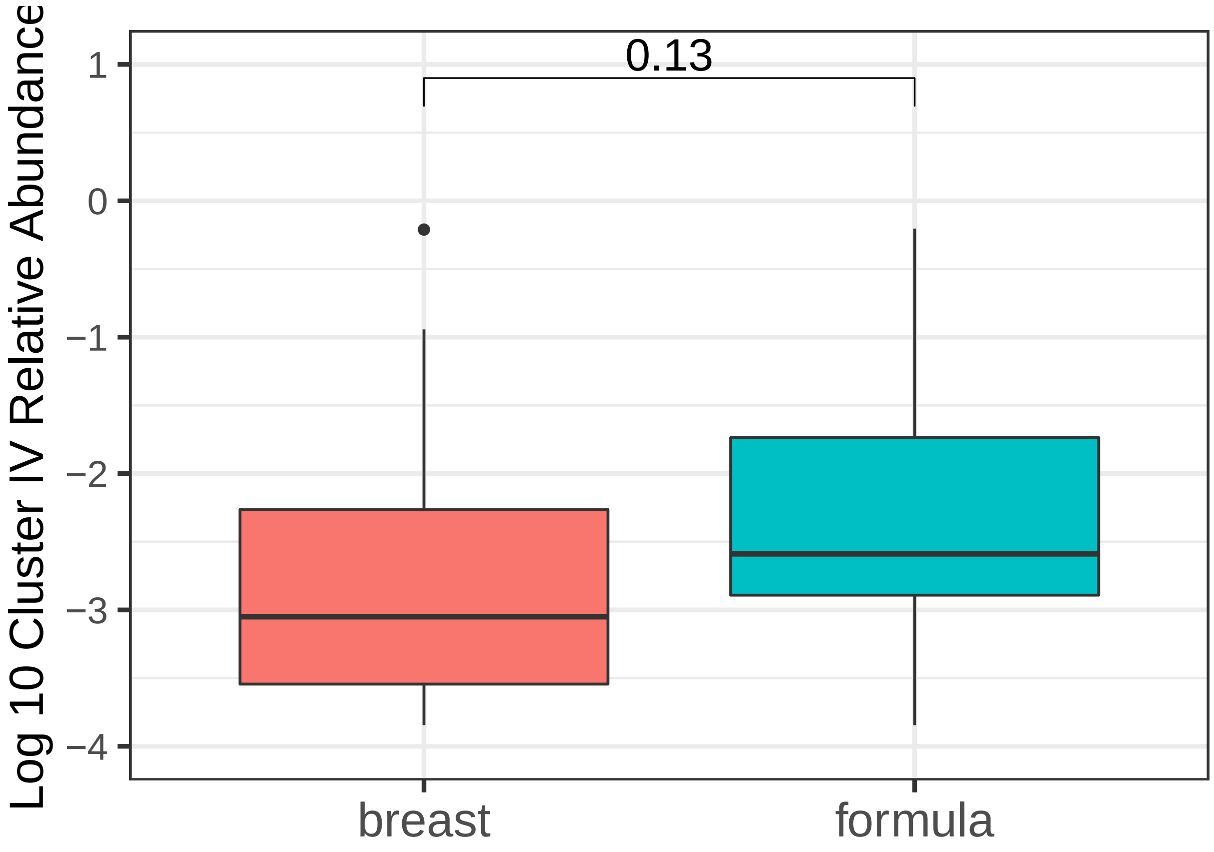


```
# ggsave("Figures/clustiv.png", plot = cluster_iv_feed, width = 4, height = 8, dpi = 600)
```

##### Correlating to Eating Behaviours

```
filtered_meta_behaviour_clust_metrics <- filtered_meta %>%
  dplyr::select(clust_i_ab,
                clust_ii_ab,
                bebq_slowness_eat,
                bebq_enjoyment_food,
                bebq_food_responsive,
                bebq_gen_appetite) %>%
  sapply(as.numeric)
```

```
## Warning in lapply(X = X, FUN = FUN, ...): NAs introduced by coercion

## Warning in lapply(X = X, FUN = FUN, ...): NAs introduced by coercion

## Warning in lapply(X = X, FUN = FUN, ...): NAs introduced by coercion

## Warning in lapply(X = X, FUN = FUN, ...): NAs introduced by coercion
```

```r
colnames(filtered_meta_behaviour_clust_metrics) <- c("Cluster I Relative Abundance", "Cluster II Relati

# Figure 3A
c <- rcorr(filtered_meta_behaviour_clust_metrics, type = "spearman")$r %>%
  round(2)
c
```

```
##                                Cluster I Relative Abundance
## Cluster I Relative Abundance                           1.00
## Cluster II Relative Abundance                         -0.25
## BEBQ Slowness of Eating                               -0.16
## BEBQ Enjoyment of Food                                 0.22
## BEBQ Food Responsiveness                              -0.23
## BEBQ General Appetite                                 -0.06
##                                Cluster II Relative Abundance
## Cluster I Relative Abundance                           -0.25
## Cluster II Relative Abundance                           1.00
## BEBQ Slowness of Eating                                 0.03
## BEBQ Enjoyment of Food                                 -0.23
## BEBQ Food Responsiveness                                0.19
## BEBQ General Appetite                                   0.12
##                                BEBQ Slowness of Eating BEBQ Enjoyment of Food
## Cluster I Relative Abundance                     -0.16                   0.22
## Cluster II Relative Abundance                     0.03                  -0.23
## BEBQ Slowness of Eating                           1.00                  -0.04
## BEBQ Enjoyment of Food                           -0.04                   1.00
## BEBQ Food Responsiveness                          0.15                  -0.04
## BEBQ General Appetite                            -0.04                   0.37
##                                BEBQ Food Responsiveness BEBQ General Appetite
## Cluster I Relative Abundance                      -0.23                 -0.06
## Cluster II Relative Abundance                      0.19                  0.12
## BEBQ Slowness of Eating                            0.15                 -0.04
## BEBQ Enjoyment of Food                            -0.04                  0.37
## BEBQ Food Responsiveness                           1.00                  0.44
## BEBQ General Appetite                              0.44                  1.00
```

```r
p <- rcorr(filtered_meta_behaviour_clust_metrics,  type = "spearman")$P %>%
  round(2)
p
```

```
##                                Cluster I Relative Abundance
## Cluster I Relative Abundance                             NA
## Cluster II Relative Abundance                          0.02
## BEBQ Slowness of Eating                                0.14
## BEBQ Enjoyment of Food                                 0.04
## BEBQ Food Responsiveness                               0.03
## BEBQ General Appetite                                  0.58
##                                Cluster II Relative Abundance
## Cluster I Relative Abundance                            0.02
## Cluster II Relative Abundance                             NA
## BEBQ Slowness of Eating                                 0.75
## BEBQ Enjoyment of Food                                  0.03
## BEBQ Food Responsiveness                                0.08
```

```
## BEBQ General Appetite                                 0.28
##                                BEBQ Slowness of Eating BEBQ Enjoyment of Food
## Cluster I Relative Abundance                      0.14                   0.04
## Cluster II Relative Abundance                     0.75                   0.03
## BEBQ Slowness of Eating                             NA                   0.72
## BEBQ Enjoyment of Food                            0.72                     NA
## BEBQ Food Responsiveness                          0.15                   0.69
## BEBQ General Appetite                             0.69                   0.00
##                                BEBQ Food Responsiveness BEBQ General Appetite
## Cluster I Relative Abundance                       0.03                  0.58
## Cluster II Relative Abundance                      0.08                  0.28
## BEBQ Slowness of Eating                            0.15                  0.69
## BEBQ Enjoyment of Food                             0.69                  0.00
## BEBQ Food Responsiveness                             NA                  0.00
## BEBQ General Appetite                              0.00                    NA
```

```r
#Get upper triangle
get_upper_tri <- function(cormat){
    cormat[lower.tri(cormat)]<- NA
    return(cormat)
  }
upper_c <- get_upper_tri(c)
upper_p <- get_upper_tri(p)

melted_c <- melt(upper_c, na.rm = TRUE)
```

```
## Warning in melt(upper_c, na.rm = TRUE): The melt generic in data.table has
## been passed a matrix and will attempt to redirect to the relevant reshape2
## method; please note that reshape2 is deprecated, and this redirection is now
## deprecated as well. To continue using melt methods from reshape2 while both
## libraries are attached, e.g. melt.list, you can prepend the namespace like
## reshape2::melt(upper_c). In the next version, this warning will become an error.
```

```r
melted_p <- melt(upper_p, na.rm = TRUE)
```

```
## Warning in melt(upper_p, na.rm = TRUE): The melt generic in data.table has
## been passed a matrix and will attempt to redirect to the relevant reshape2
## method; please note that reshape2 is deprecated, and this redirection is now
## deprecated as well. To continue using melt methods from reshape2 while both
## libraries are attached, e.g. melt.list, you can prepend the namespace like
## reshape2::melt(upper_p). In the next version, this warning will become an error.
```

```r
colnames(melted_c) <- c("Variable1", "Variable2", "Correlation")
colnames(melted_p) <- c("Variable1", "Variable2", "P")
melted_cormat <- merge(melted_c, melted_p, by=c("Variable1", "Variable2"))

# Figure 3a
ggplot(data = melted_cormat, aes(Variable2, Variable1, fill = Correlation))+
 geom_tile(color = "white")+
 scale_fill_gradient2(low = "red", high = "green",
   midpoint = 0, limit = c(-1,1), space = "Lab",
   name="Correlation Coefficient") +
```

```
  theme_minimal()+
 theme(axis.text.x = element_text(angle = 45, vjust = 1, hjust = 1))+
 coord_fixed() +
  geom_text(aes(Variable2, Variable1, label = Correlation), color = "black", size = 4) +
theme(
  axis.title.x = element_blank(),
  axis.title.y = element_blank(),
  panel.grid.major = element_blank(),
  panel.border = element_blank(),
  panel.background = element_blank(),
  axis.ticks = element_blank(),
  legend.justification = c(1, 0),
  legend.position = c(0.6, 0.8),
  legend.direction = "horizontal",
  legend.text = element_text(size = 12),
  legend.title = element_text(size = 16),
  axis.text = element_text(size = 12))+
  guides(fill = guide_colorbar(barwidth = 10, barheight = 1,
              title.position = "top", title.hjust = 0.5))
```

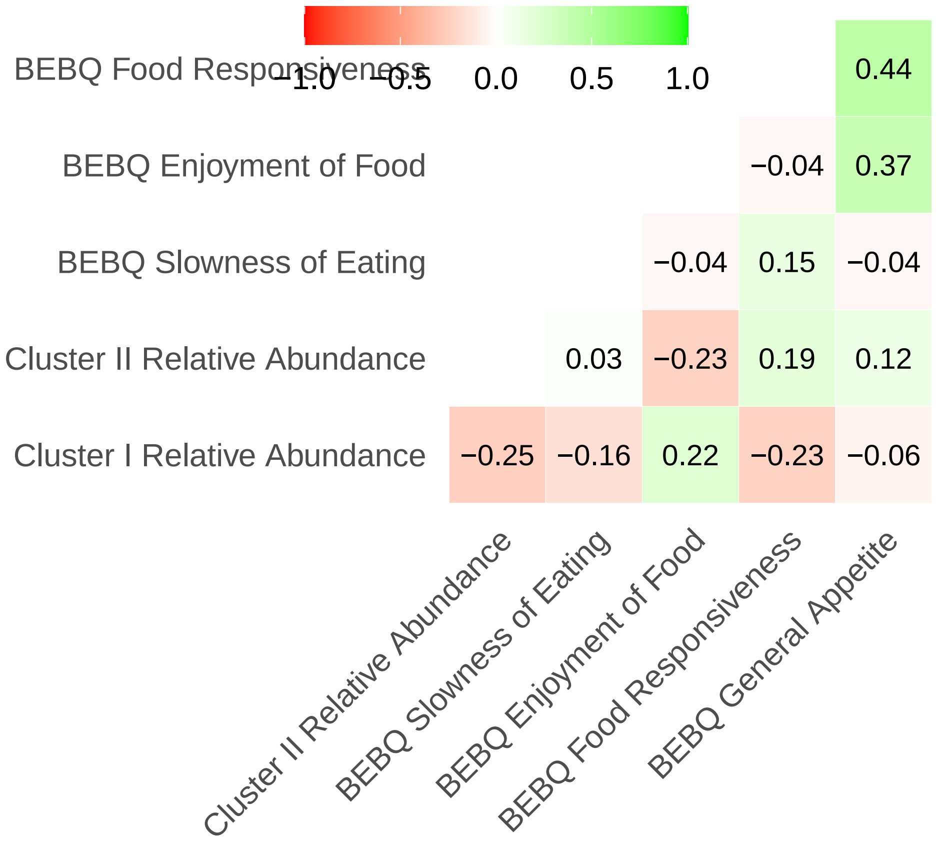


```
# ggsave("UpdatedFigs/clustbehavcor.png", width = 8, height = 8)

filtered_meta$bebq_enjoyment_food <- as.numeric(filtered_meta$bebq_enjoyment_food)
```

```
## Warning: NAs introduced by coercion
```

```r
filtered_meta$bebq_food_responsive <- as.numeric(filtered_meta$bebq_food_responsive)
```

```
## Warning: NAs introduced by coercion
```

```r
filtered_meta$bebq_gen_appetite <- as.numeric(filtered_meta$bebq_gen_appetite)
```

```
## Warning: NAs introduced by coercion
```

```r
filtered_meta$bebq_slowness_eat <- as.numeric(filtered_meta$bebq_slowness_eat)
```

```
## Warning: NAs introduced by coercion
```

```r
filtered_meta$anonymized_name <- as.factor(filtered_meta$anonymized_name)
filtered_meta <- mutate(filtered_meta,
                        log_clust_i = log10(clust_i_ab + 0.0000001),
                        log_clust_ii = log10(clust_ii_ab + 0.0000001)) #using log transformations to ma

# Figure 3b
responsive <- ggplot(aes(x = as.factor(feed), y = as.numeric(bebq_food_responsive)), data = filtered_me
  geom_boxplot(aes(fill = as.factor(feed))) +
  ylab("BEBQ Food Responsiveness") +
  theme_bw(base_size = 18) +
  xlab(NULL) +
  theme(axis.text.x = element_text(size = 18)) +
  guides(fill = FALSE) +
  ylim(0, 4.5) +
  stat_compare_means(comparisons = feed_comp, method = "wilcox.test",
                     label = "p.format", label.y.npc = 0.9, label.x = 1.3, size = 6)
responsive
```

```
## Warning: Removed 10 rows containing non-finite values (stat_boxplot).

## Warning: Removed 10 rows containing non-finite values (stat_signif).
```

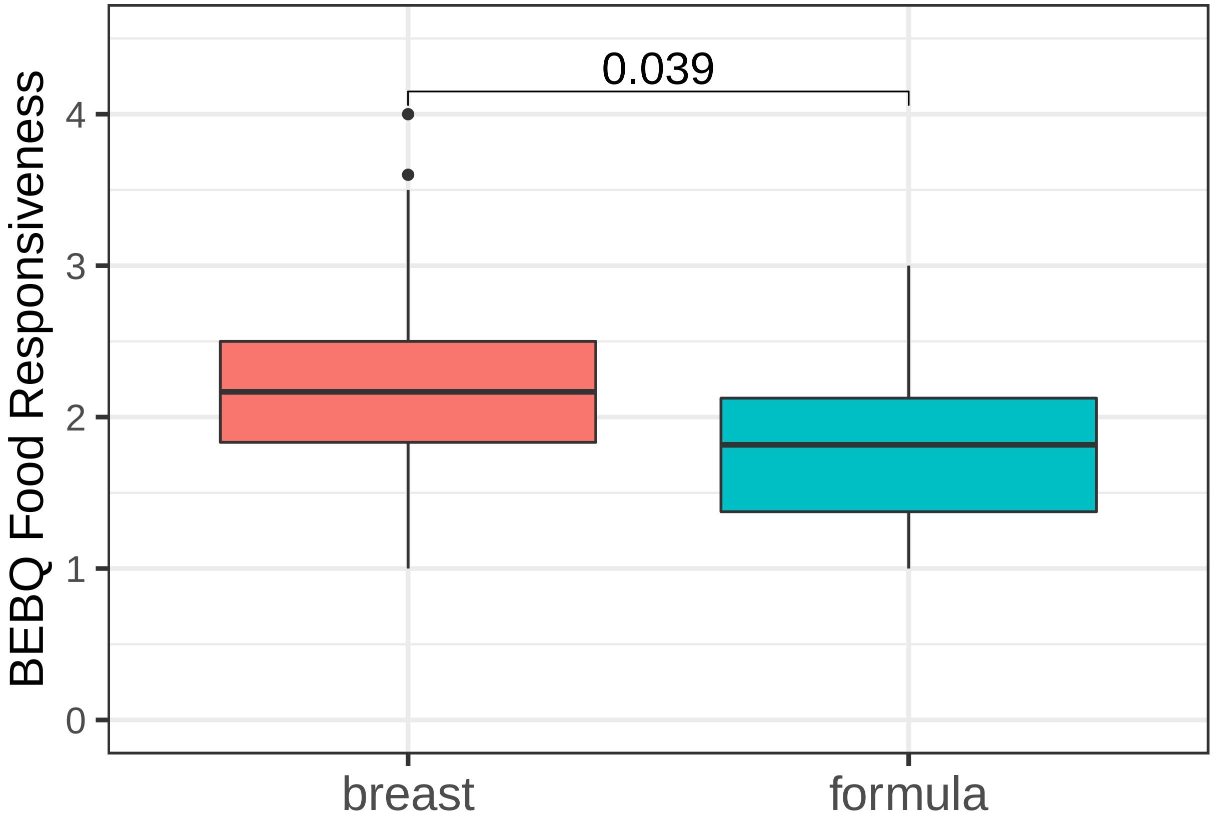


```r
# ggsave("UpdatedFigs/feedfood.png", plot = responsive, width = 6, height = 5)

# Figure 3c
clust_i_food <- ggscatter(filtered_meta, x = "bebq_food_responsive",
                              y = "log_clust_i",
                          add = "reg.line", conf.int = TRUE,
                          cor.coef = TRUE, cor.method = "spearman", cor.coef.coord = c(3, -0.5),
                          xlab = "BEBQ Food Responsiveness",
                          ylab = "Log10 Cluster I Relative Abundance") +
  theme(axis.text = element_text(size = 12),
        axis.title = element_text(size = 16))
clust_i_food
```

```
## `geom_smooth()` using formula 'y ~ x'
```

```
## Warning: Removed 10 rows containing non-finite values (stat_smooth).
```

```
## Warning: Removed 10 rows containing non-finite values (stat_cor).
```

```
## Warning: Removed 10 rows containing missing values (geom_point).
```

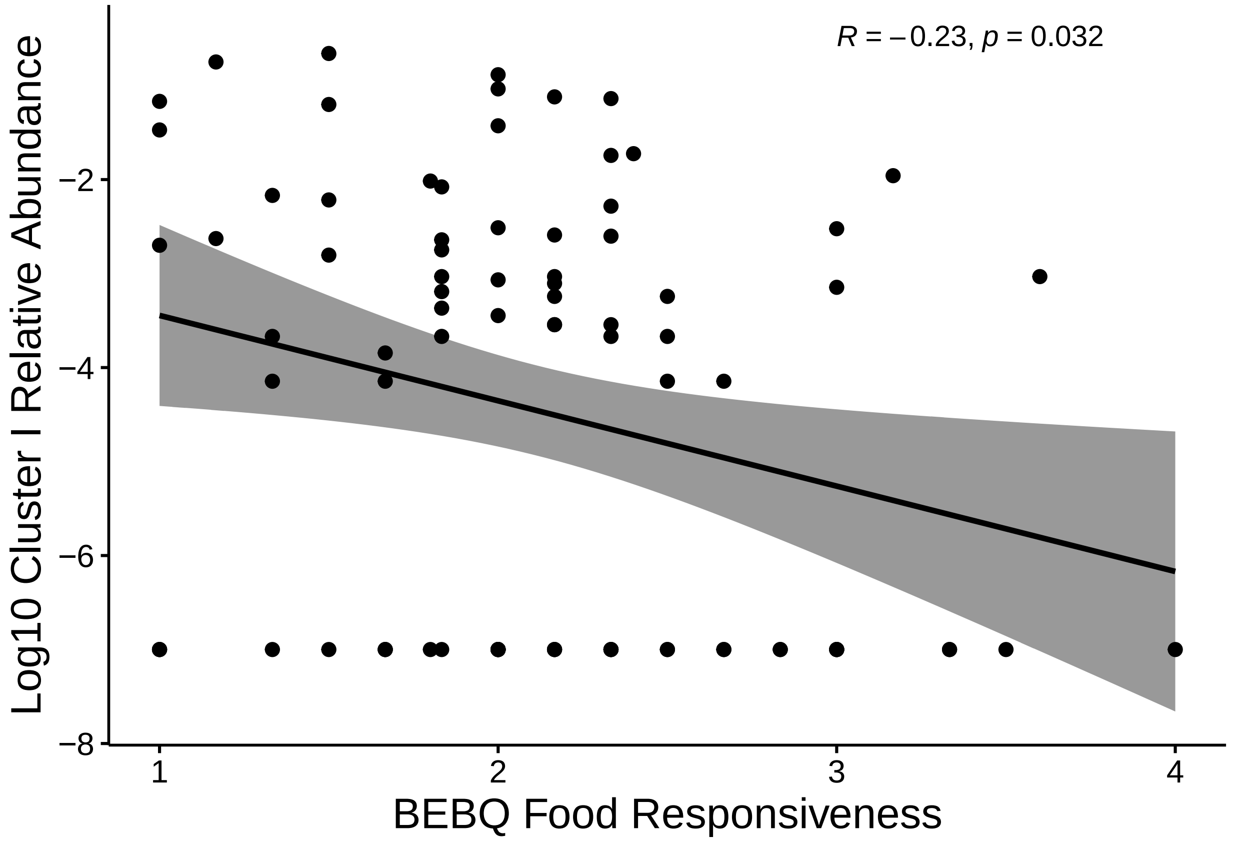


```
# ggsave("UpdatedFigs/clustifood.png", plot = clust_i_food, width = 6, height = 5)

clust_ii_food <- ggscatter(filtered_meta, x = "bebq_enjoyment_food",
                                 y = "log_clust_ii",
                              add = "reg.line", conf.int = TRUE,
                              cor.coef = TRUE, cor.method = "spearman", cor.coef.coord = c(3, -2),
                              xlab = "BEBQ Enjoyment of Food",
                              ylab = "Log10 Cluster II Relative Abundance") +
  theme(axis.text = element_text(size = 12),
        axis.title = element_text(size = 16))
clust_ii_food
```

```
## `geom_smooth()` using formula 'y ~ x'

## Warning: Removed 9 rows containing non-finite values (stat_smooth).

## Warning: Removed 9 rows containing non-finite values (stat_cor).

## Warning: Removed 9 rows containing missing values (geom_point).
```

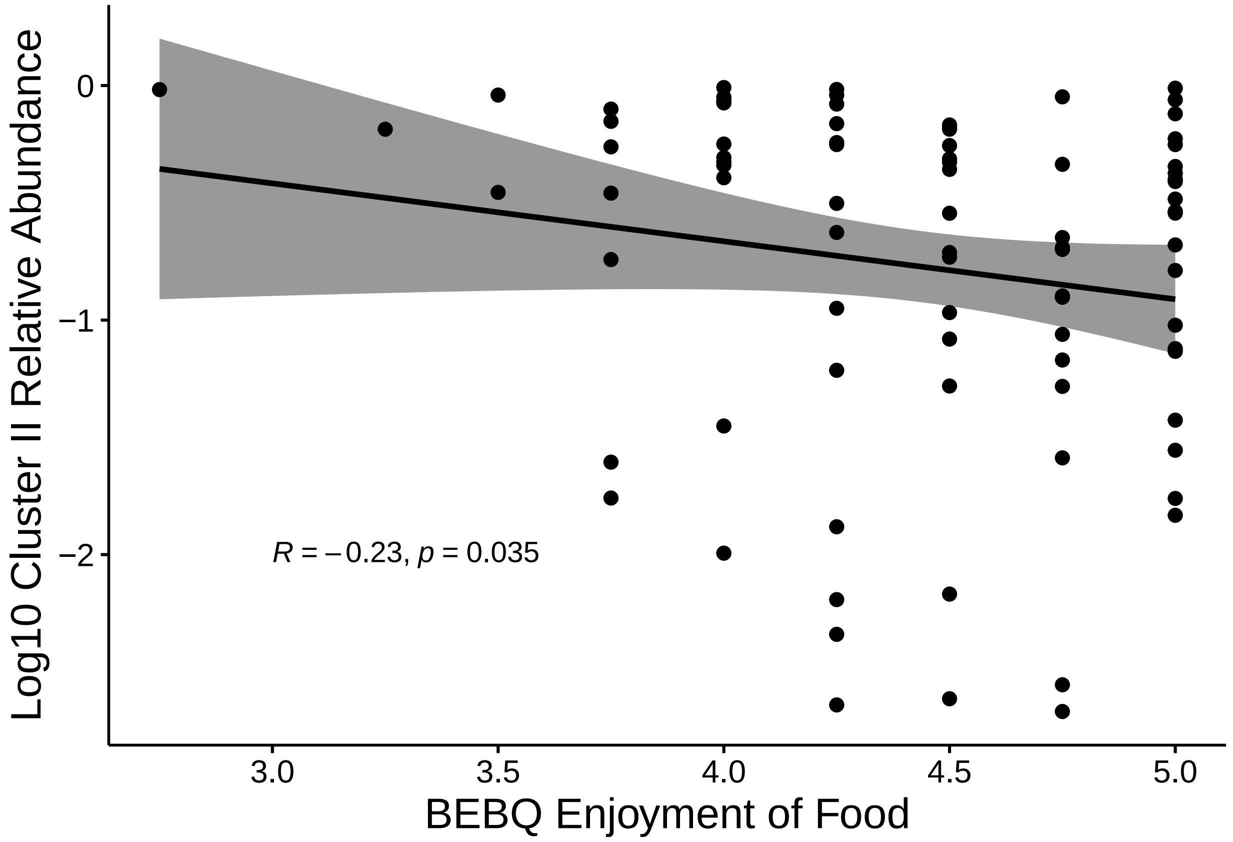


```r
# ggsave("Figures/clustiienjoy.png", plot = clust_ii_food, width = 6, height = 5)

clust_i_food <- ggscatter(filtered_meta, x = "bebq_enjoyment_food",
                                 y = "log_clust_i",
                              add = "reg.line", conf.int = TRUE,
                              cor.coef = TRUE, cor.method = "spearman", cor.coef.coord = c(3, -0.5),
                              xlab = "BEBQ Enjoyment of Food",
                              ylab = "Log10 Cluster I Relative Abundance") +
  theme(axis.text = element_text(size = 12),
        axis.title = element_text(size = 16))
clust_i_food
```

```
## `geom_smooth()` using formula 'y ~ x'

## Warning: Removed 9 rows containing non-finite values (stat_smooth).

## Warning: Removed 9 rows containing non-finite values (stat_cor).

## Warning: Removed 9 rows containing missing values (geom_point).
```

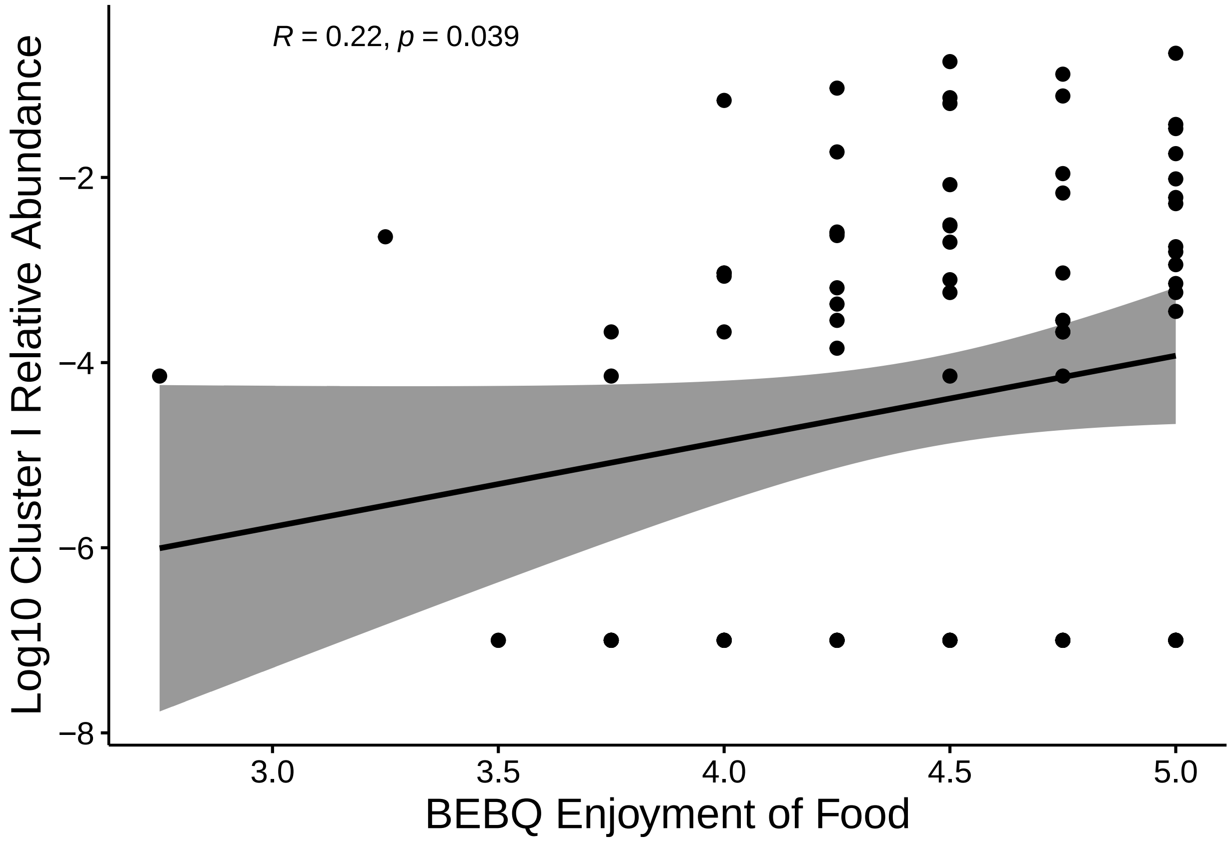


```
# ggsave("Figures/clustienjoy.png", plot = clust_i_food, width = 6, height = 5)
```

##### Pathways

```
pathway <- read_tsv("picrust/pathways_out/path_abun_unstrat_descrip.tsv")
```

```
##
## -- Column specification --------------------------------------------------------
## cols(
##   .default = col_double(),
##   pathway = col_character(),
##   description = col_character()
## )
## i Use `spec()` for the full column specifications.
```

```
pathway_10 <- pathway[rowSums(pathway == 0) >= 5, ] #filter for pathways present in at least 5% of the 

pathway_feed <- pathway_10 %>%
  dplyr::select(-pathway) %>%
  column_to_rownames(var="description") %>%
  t() %>%
  data.frame() %>%
  round() %>%
```

```r
  setDT(keep.rownames="#SampleID") %>%
  filter(`#SampleID` %in% filtered_meta$`#SampleID`) %>%
  mutate(feed = filtered_meta$feed) %>%
  dplyr::select(-feed) %>%
  column_to_rownames(var="#SampleID") %>%
  t() %>%
  as.data.frame()

conds_feed <- filtered_meta$feed
pathway.all.feed <- aldex(pathway_feed, conds_feed) %>%
  as.data.frame()
```

```
## aldex.clr: generating Monte-Carlo instances and clr values

## operating in serial mode

## computing center with all features

## aldex.ttest: doing t-test

## aldex.effect: calculating effect sizes
```

```r
pathway_sig_feed <- pathway.all.feed %>%
  filter(we.ep < 0.05, we.eBH < 0.05, wi.ep < 0.05, wi.eBH < 0.05) %>%
  setDT(keep.rownames="Pathway") %>%
  arrange()
pathway_sig_feed
```

```
##                                                     Pathway    rab.all
## 1:                              methanogenesis.from.acetate  0.7845254
## 2: L.arginine.biosynthesis.III..via.N.acetyl.L.citrulline.  9.0906583
## 3:                   acetyl.CoA.fermentation.to.butanoate.II  3.0883436
## 4:                     succinate.fermentation.to.butanoate -1.4715297
## 5:           superpathway.of.glucose.and.xylose.degradation  9.7275112
## 6:                                          tRNA.processing  9.1449637
##    rab.win.breast rab.win.formula  diff.btw diff.win     effect   overlap
## 1:      -1.220709        5.275714  5.621325 6.752128  0.8102958 0.1772576
## 2:       9.239128        8.138358 -1.088277 1.527233 -0.6996793 0.2310268
## 3:       2.714788        5.932818  3.067755 5.006555  0.6348338 0.1627648
## 4:      -2.735805        2.705530  5.359851 5.423942  0.8898627 0.1618304
## 5:       9.869060        8.657078 -1.192114 1.374338 -0.7843027 0.1852679
## 6:       9.423533        8.155247 -1.349104 1.493720 -0.8488625 0.1783724
##           we.ep      we.eBH        wi.ep      wi.eBH
## 1: 0.0004110283 0.010789776 1.232887e-04 0.007640610
## 2: 0.0016399267 0.041281059 2.043430e-03 0.041542103
## 3: 0.0032178183 0.031327357 9.525463e-05 0.007490397
## 4: 0.0008115512 0.010105690 2.109985e-04 0.009665108
## 5: 0.0002494047 0.010877293 1.831124e-04 0.008630439
## 6: 0.0001407007 0.008530787 1.711076e-04 0.008447786
```

```r
# Figure 4a
path_feed_plot <- ggplot(pathway.all.feed, aes(x = rab.all, y = diff.btw)) +
  geom_point(alpha = 0.4) +
  geom_point(data = pathway_sig_feed, aes(x = rab.all, y = diff.btw, colour = "red")) +
  xlab("Median Log2 Relative Abundance") +
  ylab("Median Log2 Difference") +
  labs(colour = NULL) +
  theme(axis.text = element_text(size = 12),
        axis.title = element_text(size = 16)) +
  annotate("text", x = c(3, -1.3), y = c(2.9, 4.9), label = c("Acetyl-CoA", "Succinate"))
path_feed_plot
```

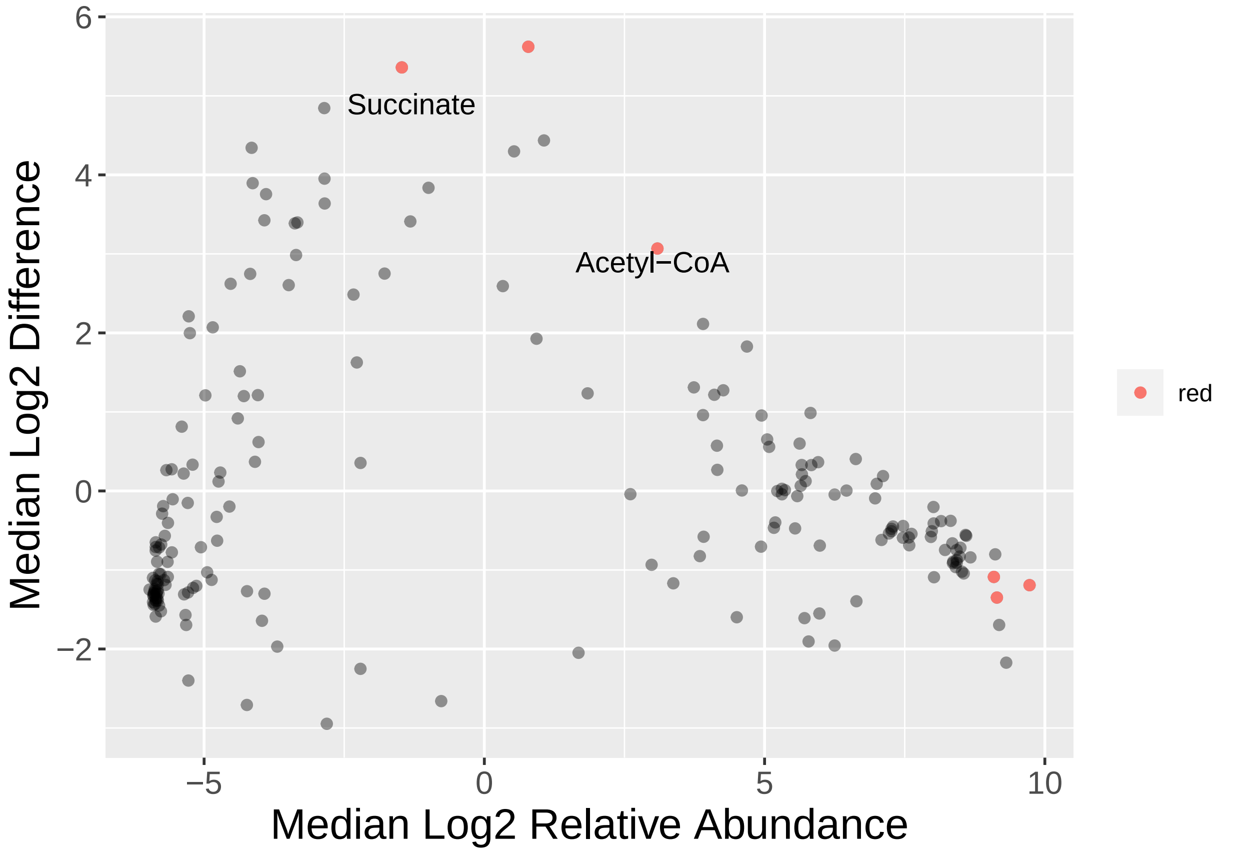


```r
# ggsave("UpdatedFigs/pathways.png", plot = path_feed_plot, width = 6, height = 4)
```

##### Correlation to Feed and Eating Behaviours

```r
rel_ab <- function(x) x/sum(x)

pathway_spectrum <- pathway %>%
  dplyr::select(-pathway) %>%
  column_to_rownames(var="description") %>%
  rel_ab() %>%
  t() %>%
```

```
  data.frame() %>%
  setDT(keep.rownames="#SampleID") %>%
  filter(`#SampleID` %in% filtered_meta$`#SampleID`) %>%
  mutate(bebq_enjoyment_food = filtered_meta$bebq_enjoyment_food,
         bebq_food_responsive = filtered_meta$bebq_food_responsive,
         bebq_slowness_eat = filtered_meta$bebq_slowness_eat,
         bebq_gen_appetite = filtered_meta$bebq_gen_appetite,
         log_clust_i = filtered_meta$log_clust_i,
         log_clust_ii = filtered_meta$log_clust_ii) %>%
  mutate(log_acet = log10(acetyl.CoA.fermentation.to.butanoate.II + 0.0000001),
         log_succ = log10(succinate.fermentation.to.butanoate + 0.0000001)) %>%
  as.data.frame() %>%
  sapply(as.numeric)
```

```
## Warning in lapply(X = X, FUN = FUN, ...): NAs introduced by coercion
```

```
pathway_spec_feed <- pathway_spectrum %>%
  as.data.frame() %>%
  mutate(feed = filtered_meta$feed) %>%
  as.data.frame()

acet <- ggplot(aes(x = as.factor(feed), y = log_acet), data = pathway_spec_feed) +
  geom_boxplot(aes(fill = as.factor(feed))) +
  ylab(expression(atop("Log10 Acetyl-CoA Fermentation to Butanoate II", paste("(Relative Abundance)"))))
  theme_bw(base_size = 18) +
  xlab(NULL) +
  theme(axis.text.x = element_text(size = 18)) +
  guides(fill = FALSE) +
  ylim(-7, -2.5) +
  stat_compare_means(comparisons = feed_comp, method = "wilcox.test",
                     label = "p.format", label.y.npc = 0.9, label.x = 1.3, size = 6)
acet
```

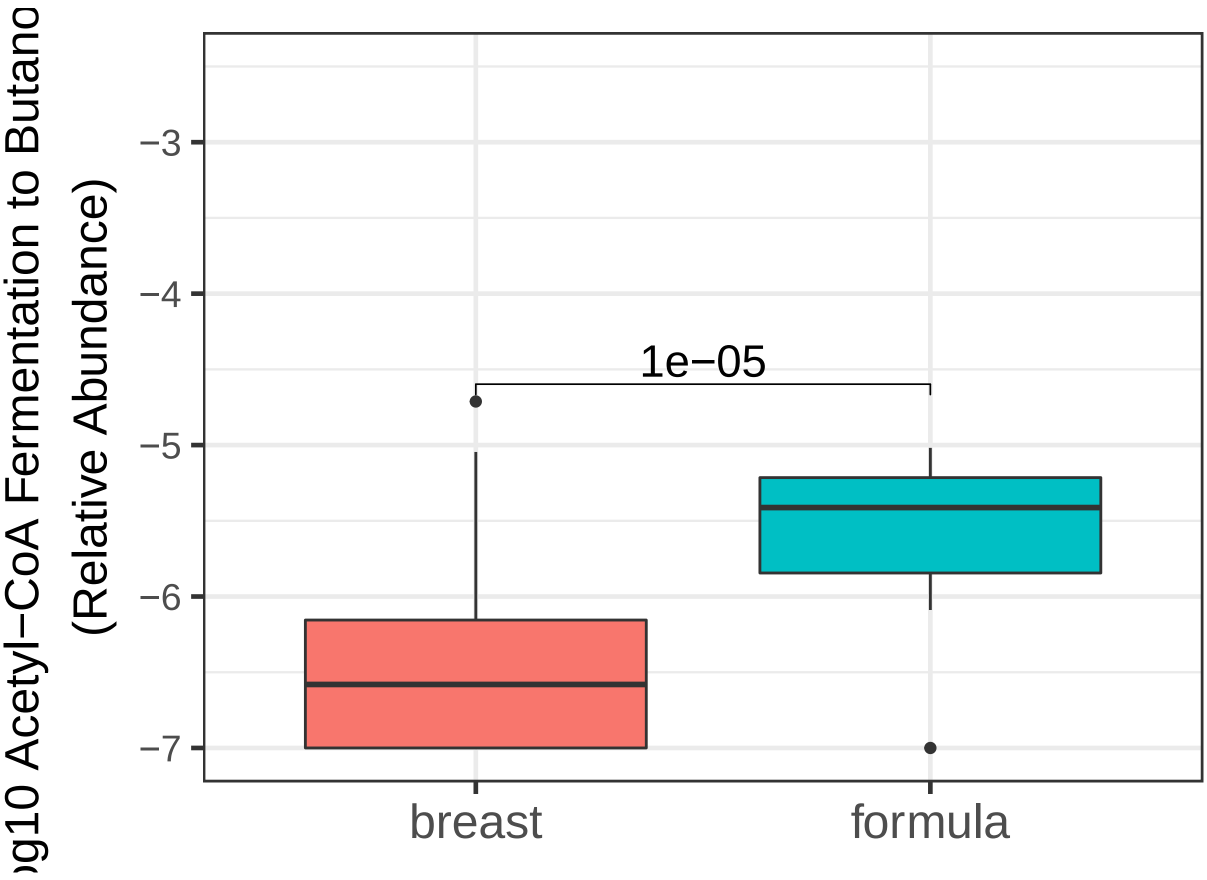


```r
# ggsave("Figures/acetfeed.png", plot = acet, width = 4, height = 8)

# Figure 4b
succ <- ggplot(aes(x = as.factor(feed), y = log_succ), data = pathway_spec_feed) +
  geom_boxplot(aes(fill = as.factor(feed))) +
  ylab(expression(atop("Log10 Succinate Fermentation to Butanoate", paste("(Relative Abundance)")))) +
  theme_bw(base_size = 18) +
  xlab(NULL) +
  theme(axis.text.x = element_text(size = 18)) +
  guides(fill = FALSE) +
  ylim(-7, -5) +
  stat_compare_means(comparisons = feed_comp, method = "wilcox.test",
                     label = "p.format", label.y.npc = 0.9, label.x = 1.3, size = 6)
succ
```

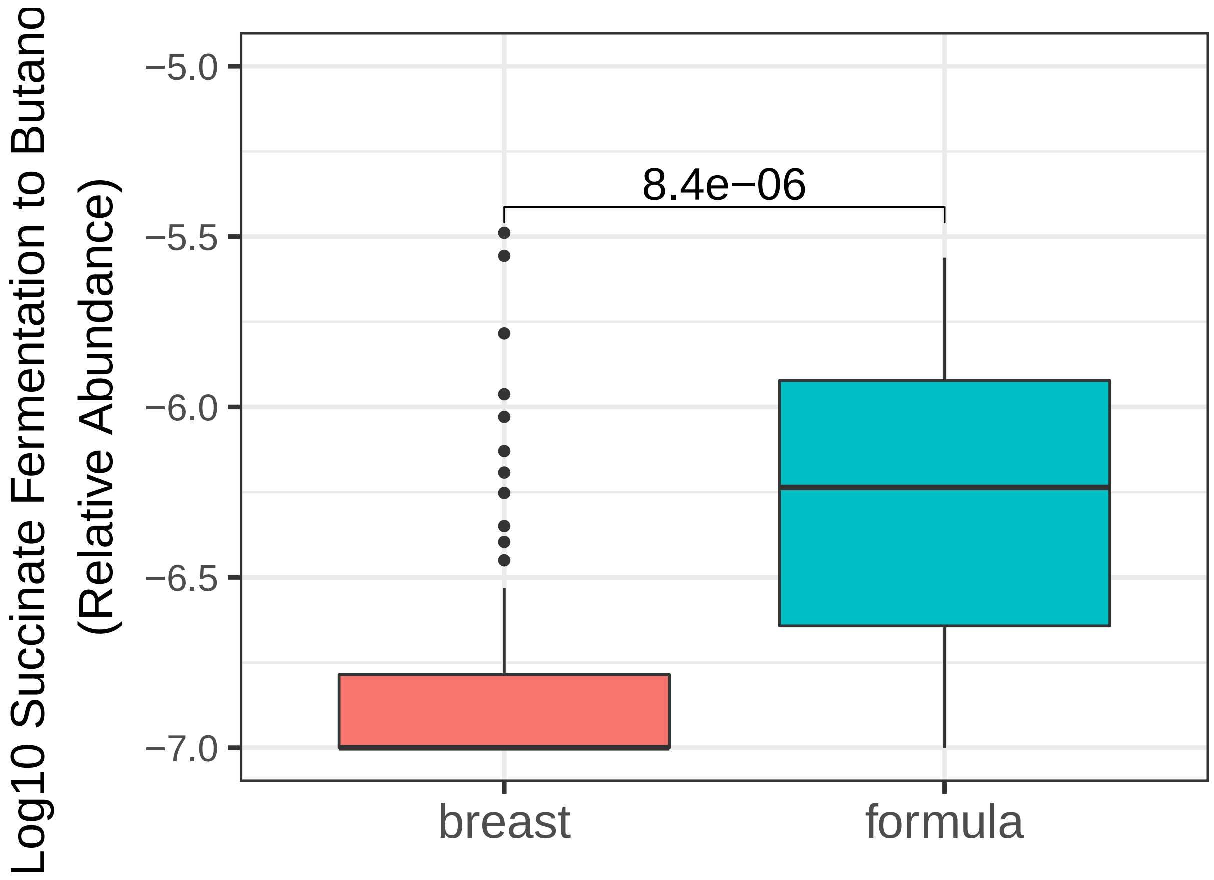


```
# ggsave("UpdatedFigs/succfeed.png", plot = succ, width = 4, height = 8)

pathway_correlation <- pathway_spectrum %>%
  as.data.frame() %>%
  dplyr::select(bebq_enjoyment_food,
         bebq_food_responsive,
         bebq_slowness_eat,
         bebq_gen_appetite,
         log_clust_i,
         log_clust_ii,
         acetyl.CoA.fermentation.to.butanoate.II,
         succinate.fermentation.to.butanoate) %>%
  sapply(as.numeric)

colnames(pathway_correlation) <- c("BEBQ Enjoyment of Food", "BEBQ Food Responsiveness","BEBQ Slowness o

# Figure 4E
c <- rcorr(pathway_correlation, type = "spearman")$r %>%
  round(2)
c
```

```
##                              BEBQ Enjoyment of Food BEBQ Food Responsiveness
## BEBQ Enjoyment of Food                         1.00                    -0.04
## BEBQ Food Responsiveness                      -0.04                     1.00
## BEBQ Slowness of Eating                       -0.04                     0.15
## BEBQ General Appetite                          0.37                     0.44
```

```
## Cluster I Relative Abundance                    0.22                   -0.23
## Cluster II Relative Abundance                  -0.23                    0.19
## Acetyl-CoA                                      0.21                   -0.06
## Succinate                                       0.12                   -0.23
##                               BEBQ Slowness of Eating BEBQ General Appetite
## BEBQ Enjoyment of Food                          -0.04                  0.37
## BEBQ Food Responsiveness                         0.15                  0.44
## BEBQ Slowness of Eating                          1.00                 -0.04
## BEBQ General Appetite                           -0.04                  1.00
## Cluster I Relative Abundance                    -0.16                 -0.06
## Cluster II Relative Abundance                    0.03                  0.12
## Acetyl-CoA                                      -0.38                  0.09
## Succinate                                       -0.38                 -0.05
##                               Cluster I Relative Abundance
## BEBQ Enjoyment of Food                                0.22
## BEBQ Food Responsiveness                             -0.23
## BEBQ Slowness of Eating                              -0.16
## BEBQ General Appetite                                -0.06
## Cluster I Relative Abundance                          1.00
## Cluster II Relative Abundance                        -0.25
## Acetyl-CoA                                            0.54
## Succinate                                             0.67
##                               Cluster II Relative Abundance Acetyl-CoA
## BEBQ Enjoyment of Food                                -0.23       0.21
## BEBQ Food Responsiveness                               0.19      -0.06
## BEBQ Slowness of Eating                                0.03      -0.38
## BEBQ General Appetite                                  0.12       0.09
## Cluster I Relative Abundance                          -0.25       0.54
## Cluster II Relative Abundance                          1.00      -0.19
## Acetyl-CoA                                            -0.19       1.00
## Succinate                                             -0.08       0.72
##                               Succinate
## BEBQ Enjoyment of Food             0.12
## BEBQ Food Responsiveness          -0.23
## BEBQ Slowness of Eating           -0.38
## BEBQ General Appetite             -0.05
## Cluster I Relative Abundance       0.67
## Cluster II Relative Abundance     -0.08
## Acetyl-CoA                         0.72
## Succinate                          1.00
```

```
p <- rcorr(pathway_correlation,  type = "spearman")$P %>%
  round(2)
p
```

```
##                               BEBQ Enjoyment of Food BEBQ Food Responsiveness
## BEBQ Enjoyment of Food                            NA                     0.69
## BEBQ Food Responsiveness                        0.69                       NA
## BEBQ Slowness of Eating                         0.72                     0.15
## BEBQ General Appetite                           0.00                     0.00
## Cluster I Relative Abundance                    0.04                     0.03
## Cluster II Relative Abundance                   0.03                     0.08
## Acetyl-CoA                                      0.05                     0.56
## Succinate                                       0.26                     0.04
```

```
##                                BEBQ Slowness of Eating BEBQ General Appetite
## BEBQ Enjoyment of Food                            0.72                  0.00
## BEBQ Food Responsiveness                          0.15                  0.00
## BEBQ Slowness of Eating                             NA                  0.69
## BEBQ General Appetite                             0.69                    NA
## Cluster I Relative Abundance                      0.14                  0.58
## Cluster II Relative Abundance                     0.75                  0.28
## Acetyl-CoA                                        0.00                  0.42
## Succinate                                         0.00                  0.68
##                                Cluster I Relative Abundance
## BEBQ Enjoyment of Food                                 0.04
## BEBQ Food Responsiveness                               0.03
## BEBQ Slowness of Eating                                0.14
## BEBQ General Appetite                                  0.58
## Cluster I Relative Abundance                             NA
## Cluster II Relative Abundance                          0.02
## Acetyl-CoA                                             0.00
## Succinate                                              0.00
##                                Cluster II Relative Abundance Acetyl-CoA
## BEBQ Enjoyment of Food                                  0.03       0.05
## BEBQ Food Responsiveness                                0.08       0.56
## BEBQ Slowness of Eating                                 0.75       0.00
## BEBQ General Appetite                                   0.28       0.42
## Cluster I Relative Abundance                            0.02       0.00
## Cluster II Relative Abundance                             NA       0.07
## Acetyl-CoA                                              0.07         NA
## Succinate                                               0.41       0.00
##                                Succinate
## BEBQ Enjoyment of Food              0.26
## BEBQ Food Responsiveness            0.04
## BEBQ Slowness of Eating             0.00
## BEBQ General Appetite               0.68
## Cluster I Relative Abundance        0.00
## Cluster II Relative Abundance       0.41
## Acetyl-CoA                          0.00
## Succinate                             NA
```

```
upper_c <- get_upper_tri(c)
upper_p <- get_upper_tri(p)

melted_c <- melt(upper_c, na.rm = TRUE)
```

```
## Warning in melt(upper_c, na.rm = TRUE): The melt generic in data.table has
## been passed a matrix and will attempt to redirect to the relevant reshape2
## method; please note that reshape2 is deprecated, and this redirection is now
## deprecated as well. To continue using melt methods from reshape2 while both
## libraries are attached, e.g. melt.list, you can prepend the namespace like
## reshape2::melt(upper_c). In the next version, this warning will become an error.
```

```
melted_p <- melt(upper_p, na.rm = TRUE)
```

```
## Warning in melt(upper_p, na.rm = TRUE): The melt generic in data.table has
## been passed a matrix and will attempt to redirect to the relevant reshape2
```

```
## method; please note that reshape2 is deprecated, and this redirection is now
## deprecated as well. To continue using melt methods from reshape2 while both
## libraries are attached, e.g. melt.list, you can prepend the namespace like
## reshape2::melt(upper_p). In the next version, this warning will become an error.
```

```
colnames(melted_c) <- c("Variable1", "Variable2", "Correlation")
colnames(melted_p) <- c("Variable1", "Variable2", "P")
melted_cormat <- merge(melted_c, melted_p, by=c("Variable1", "Variable2"))

ggplot(data = melted_cormat, aes(Variable2, Variable1, fill = Correlation))+
 geom_tile(color = "white")+
 scale_fill_gradient2(low = "red", high = "green",
   midpoint = 0, limit = c(-1,1), space = "Lab",
   name="Correlation Coefficient") +
  theme_minimal()+
 theme(axis.text.x = element_text(angle = 45, vjust = 1, hjust = 1))+
 coord_fixed() +
  geom_text(aes(Variable2, Variable1, label = Correlation), color = "black", size = 4) +
theme(
  axis.title.x = element_blank(),
  axis.title.y = element_blank(),
  panel.grid.major = element_blank(),
  panel.border = element_blank(),
  panel.background = element_blank(),
  axis.ticks = element_blank(),
  legend.justification = c(1, 0),
  legend.position = c(0.6, 0.8),
  legend.direction = "horizontal",
  legend.text = element_text(size = 12),
  legend.title = element_text(size = 16),
  axis.text = element_text(size = 12))+
  guides(fill = guide_colorbar(barwidth = 10, barheight = 1,
                title.position = "top", title.hjust = 0.5))
```

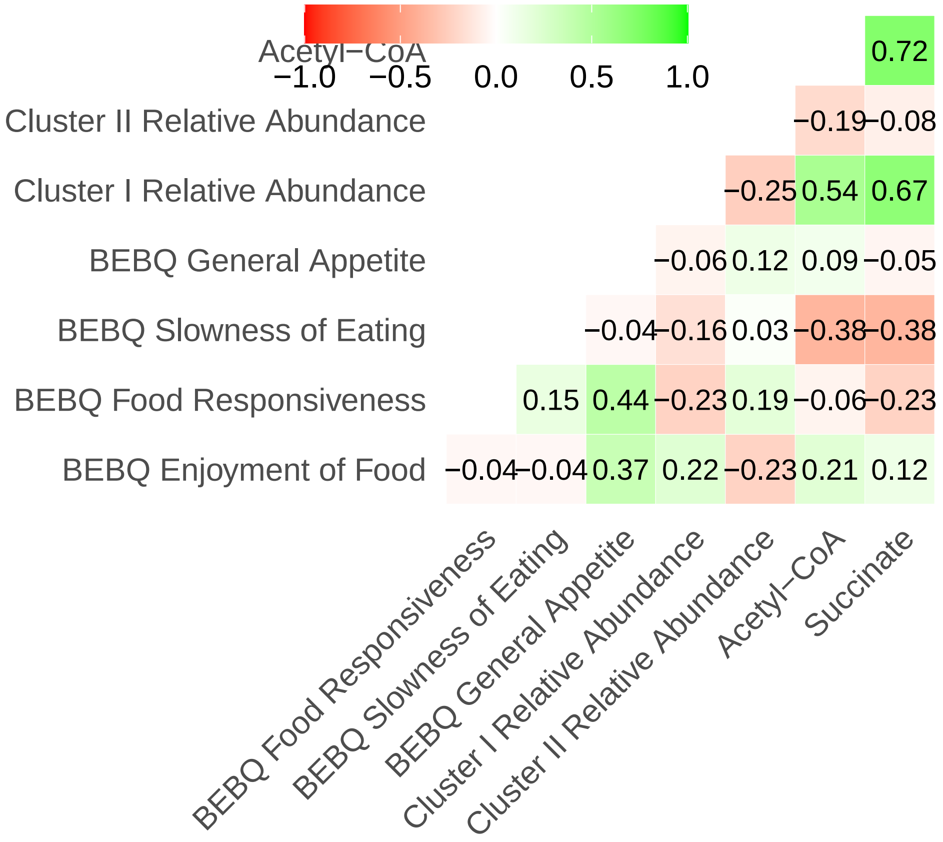


```
# ggsave("Figures/pathfeed.png", width = 8, height = 8)
```

**Correlation to Cluster I and Food Responsiveness**

```
# Figure 4c
succ_clust_i <- ggscatter(pathway_spec_feed, x = "log_clust_i",
                                 y = "log_succ",
                              add = "reg.line", conf.int = TRUE,
                              cor.coef = TRUE, cor.method = "spearman",
                              xlab = "Log 10 Cluster I Relative Abundance",
                              ylab = (expression(atop("Log10 Succinate Fermentation to Butanoate", paste
  theme(axis.text = element_text(size = 12),
        axis.title = element_text(size = 16))
succ_clust_i
```

```
## `geom_smooth()` using formula 'y ~ x'
```

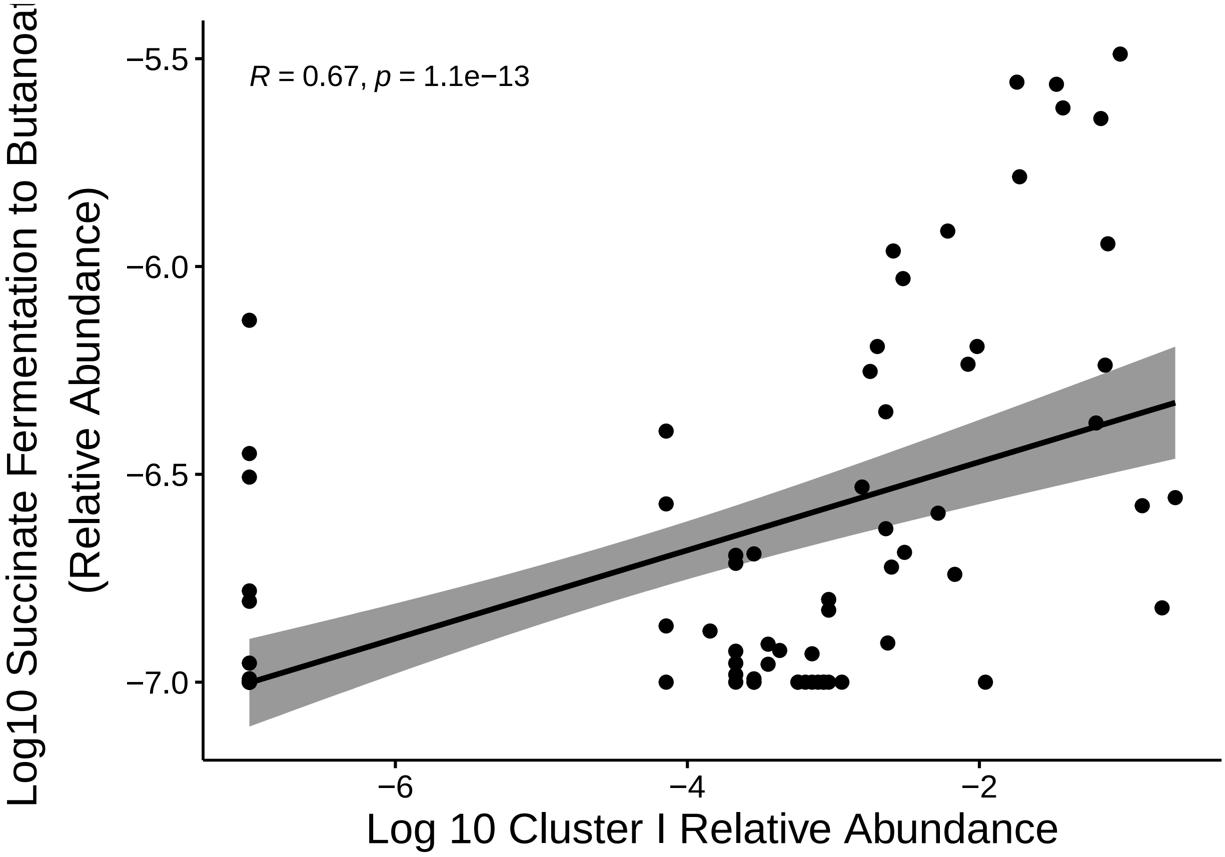


```r
# ggsave("UpdatedFigs/succclusti.png", plot = succ_clust_i, width = 6, height = 5, units = "in")

# Figure 4d
succ_food <- ggscatter(pathway_spec_feed, x = "log_succ",
                              y = "bebq_food_responsive",
                        add = "reg.line", conf.int = TRUE,
                        cor.coef = TRUE, cor.method = "spearman",
                        ylab = "BEBQ Food Responsiveness",
                        xlab = (expression(atop("Log10 Succinate Fermentation to Butanoate", paste
  theme(axis.text = element_text(size = 12),
        axis.title = element_text(size = 16))
succ_food
```

```
## `geom_smooth()` using formula 'y ~ x'

## Warning: Removed 10 rows containing non-finite values (stat_smooth).

## Warning: Removed 10 rows containing non-finite values (stat_cor).

## Warning: Removed 10 rows containing missing values (geom_point).
```

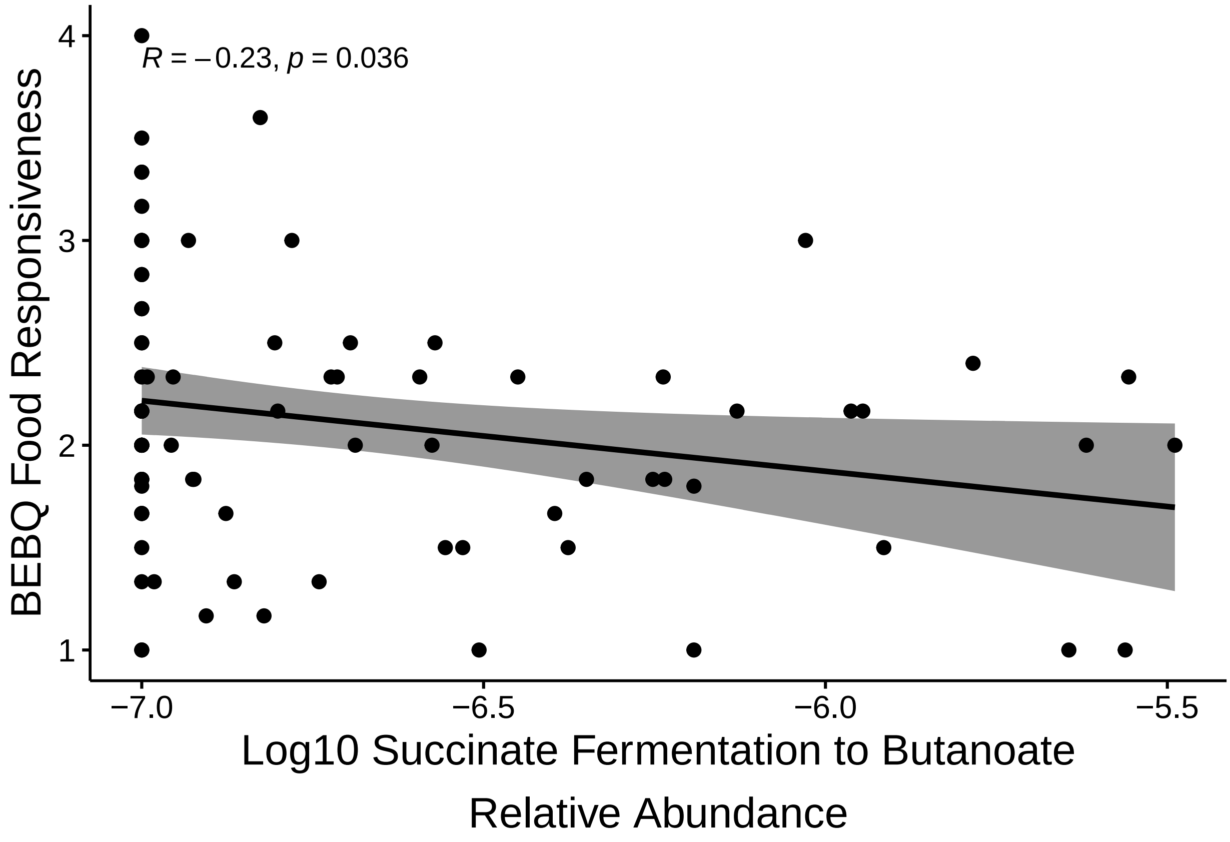


```
# ggsave("UpdatedFigs/succfood.png", plot = succ_food, width = 6, height = 5, units = "in")
```
