## Supplemental Tables and Figures for "Diet Potentially Drives the Differentiation of Eating Behaviours via Alterations to the Gut Microbiome in Infants"

**Contents:**

**Supplementary Table 1 Top ten most important genera for predicting diet belong to the phylum Firmicutes.** Variable importance table for random forest classifier trained to predict diet based on genus relative abundance.

**Supplementary Table 2 Two out of the six significantly differentially abundant pathways are related to SCFA synthesis**. Pathways were inferred using PICRUSt2 and differential abundance was analyzed using ALDeX2. SCFA synthesis pathways are underlined.

**Supplementary Fig. 1 Beta diversity distances for breastfed and formula-fed infants.** PCoA plots for (A) unweighted UniFrac, (B) Bray-Curtis, and (C) Jaccard. Each point represents an infant included in this study and is coloured by diet.

**Supplementary Fig. 2 The optimal sequencing depth is 14000.** (A) Alpha rarefaction plot for infant samples coloured by diet. (B) The number of infant samples retained at each sequencing depth coloured by diet. (C) Histogram showing the distribution of all samples across sequencing depths.

**Supplementary Fig. 3 Gut microbiomes of formula-fed infants consistently have higher alpha diversity than those of breastfed infants.** Boxplots for observed OTUs, Chao1, ACE, Shannon, Simpson, Inverse Simpson, and Fisher alpha diversity metrics separated by feed.

**Supplementary Fig. 4 Clusters III and IV are not differentially abundant between breastfed and formula-fed infants**. (A) Heatmap based on the covariance of bacterial genera coloured by Spearman correlation coefficient. Axes are arranged based on Spearman correlation distance and Ward linkage. (B) Comparison of cluster III and IV relative abundances across feeding groups using the Mann-Whitney U test.

**Supplementary Fig. 5 BEBQ enjoyment of food is also significantly correlated with cluster abundances.** (A-B) Spearman correlation coefficient and significance for clusters I and II and BEBQ enjoyment of food.

**Supplementary Fig. 6 Pathway relative abundance differs significantly between breastfed and formula-fed infants, and is significantly correlated with cluster I relative abundance.** (A) Acetyl-CoA fermentation to butanoate II is significantly more highly expressed in formula-fed infants according to the Mann-Whitney U test. (B) Heatmap of correlation matrix between cluster abundance, food responsiveness, and SCFA pathways coloured by Spearman correlation coefficient. Significance ($p < 0.05$) is marked with a star (*).

**Supplementary Table 1 Top ten most important genera for predicting diet belong to the phylum Firmicutes.** Variable importance table for random forest classifier trained to predict diet based on genus relative abundance.

| **Genus** | **Importance** |
|---|---|
| *Blautia* | 100.00 |
| *Eubacterium* | 80.09 |
| *Dorea* | 59.12 |
| *Akkermansia* | 56.58 |
| *Eggerthella* | 53.22 |
| *Haemophilus* | 49.08 |
| *Coprococcus* | 48.04 |
| *Acinetobacter* | 46.38 |
| *Proteus* | 44.50 |
| *Ruminococcus* | 43.07 |

**Supplementary Table 2 Two out of the six significantly differentially abundant pathways are related to SCFA synthesis**. Pathways were inferred using PICRUSt2 and differential abundance was analyzed using ALDeX2. SCFA synthesis pathways are underlined.

| **Pathway** | **Welch's t-test (corrected P-value)** | **Wilcoxon test (corrected P-value)** |
|---|---|---|
| Methanogenesis from acetate | 0.011 | 0.0076 |
| L-arginine biosynthesis III via N-acetyl L-citrulline. | 0.041 | 0.042 |
| <u>Acetyl-CoA fermentation to butanoate II</u> | 0.031 | 0.0075 |
| <u>Succinate fermentation to butanoate</u> | 0.010 | 0.0097 |
| Superpathway of glucose and xylose degradation | 0.011 | 0.0086 |
| tRNA processing | 0.0085 | 0.0085 |

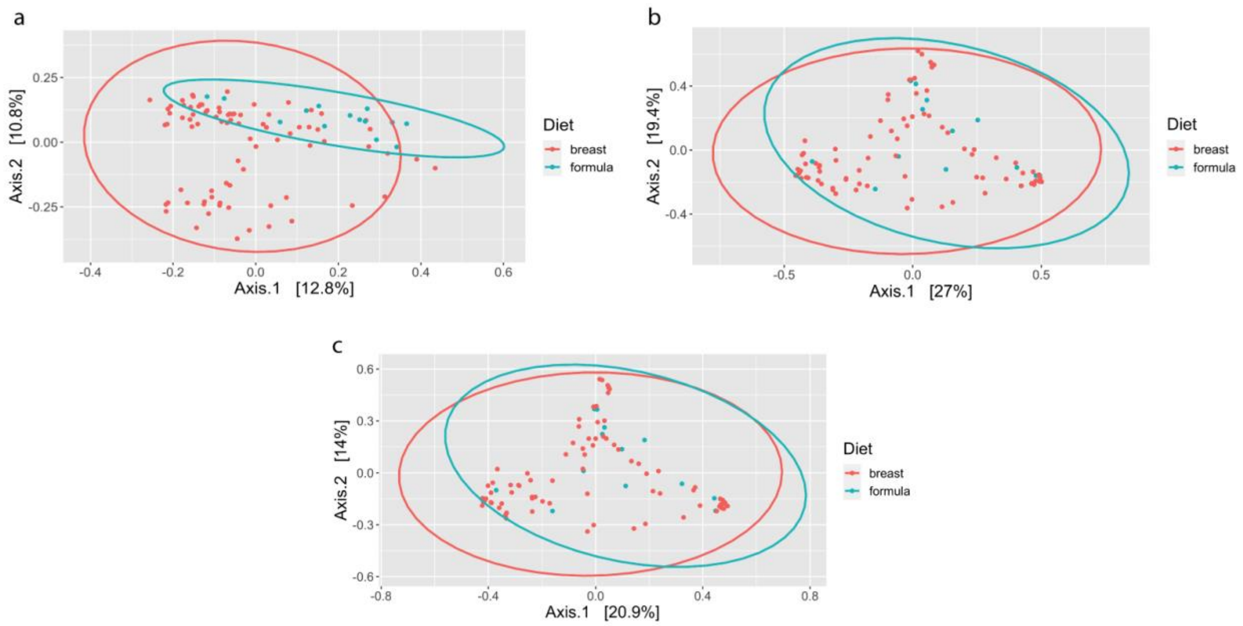


**Supplementary Fig. 1 Beta diversity distances for breastfed and formula-fed infants.** PCoA plots for (A) unweighted UniFrac, (B) Bray-Curtis, and (C) Jaccard. Each point represents an infant included in this study and is coloured by diet.

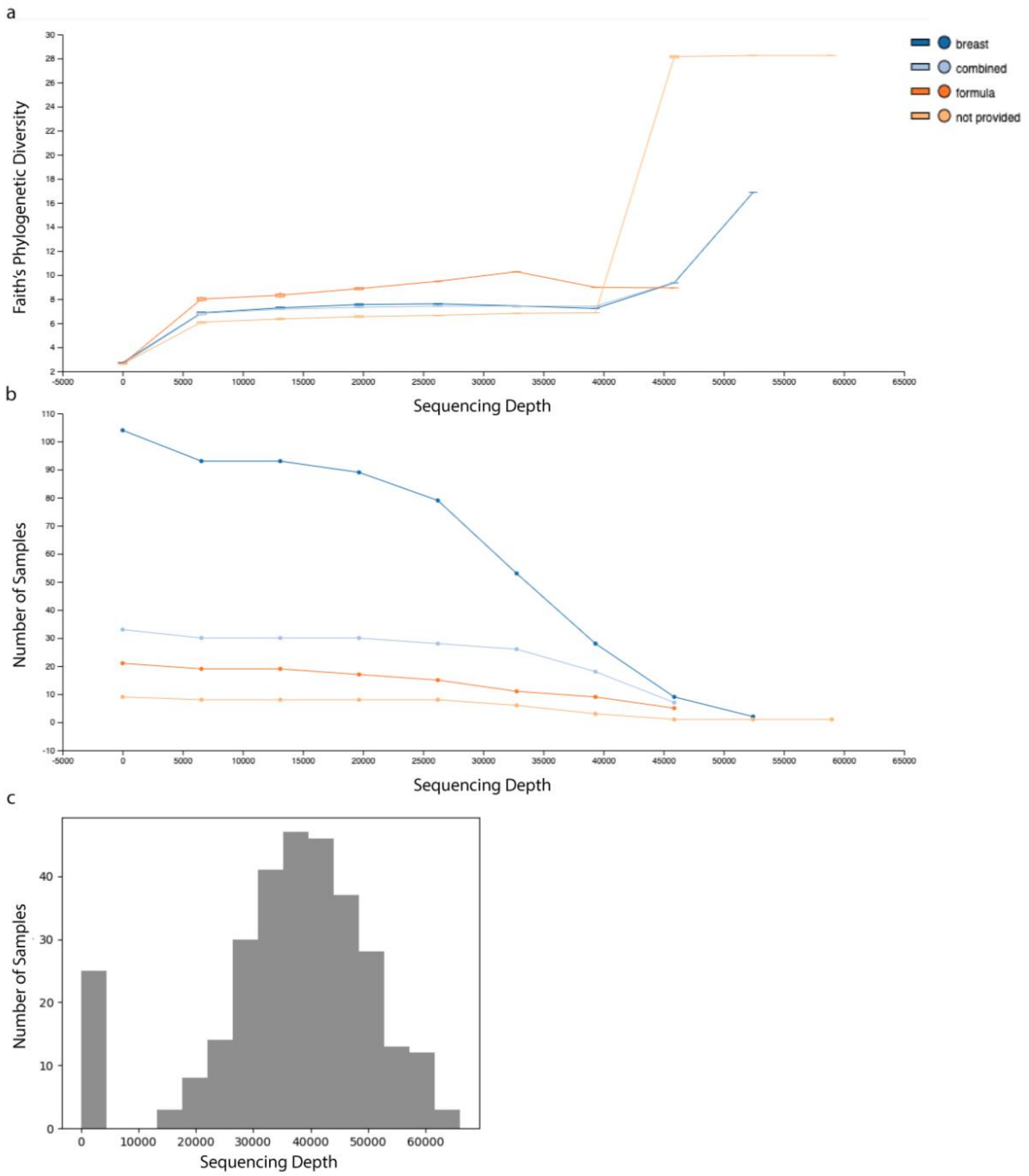


**Supplementary Fig. 2 The optimal sequencing depth is 14000.** (A) Alpha rarefaction plot for infant samples coloured by diet. (B) The number of infant samples retained at each sequencing depth coloured by diet. (C) Histogram showing the distribution of all samples across sequencing depths.

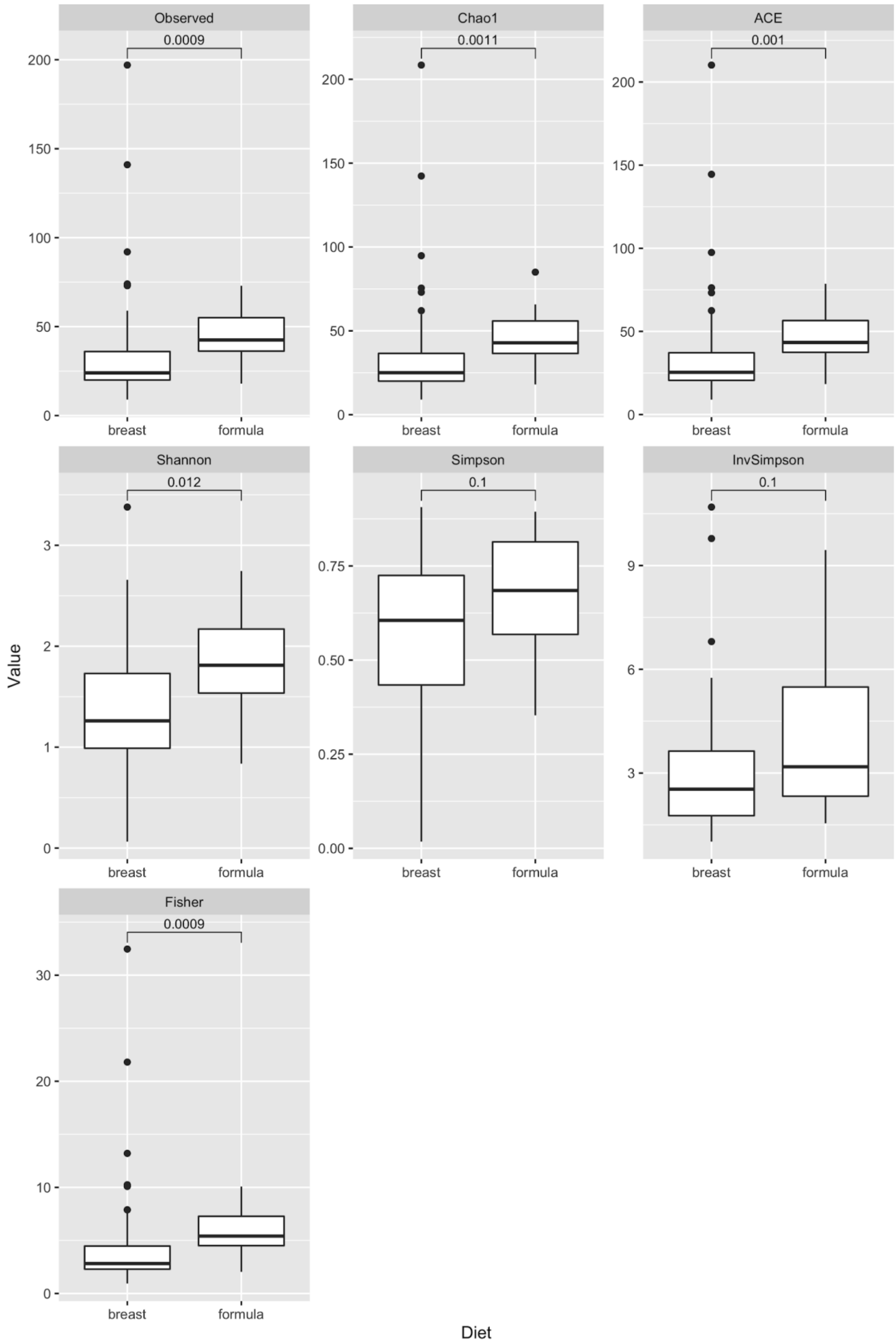

Observed
0.0009
Chao1
0.0011
ACE
0.001
Shannon
0.012
Simpson
0.1
InvSimpson
0.1
Fisher
0.0009
breast
formula
Value
Diet

**Supplementary Fig. 3 Gut microbiomes of formula-fed infants consistently have higher alpha diversity than those of breastfed infants.** Boxplots for observed OTUs, Chao1, ACE, Shannon, Simpson, Inverse Simpson, and Fisher alpha diversity metrics separated by diet.

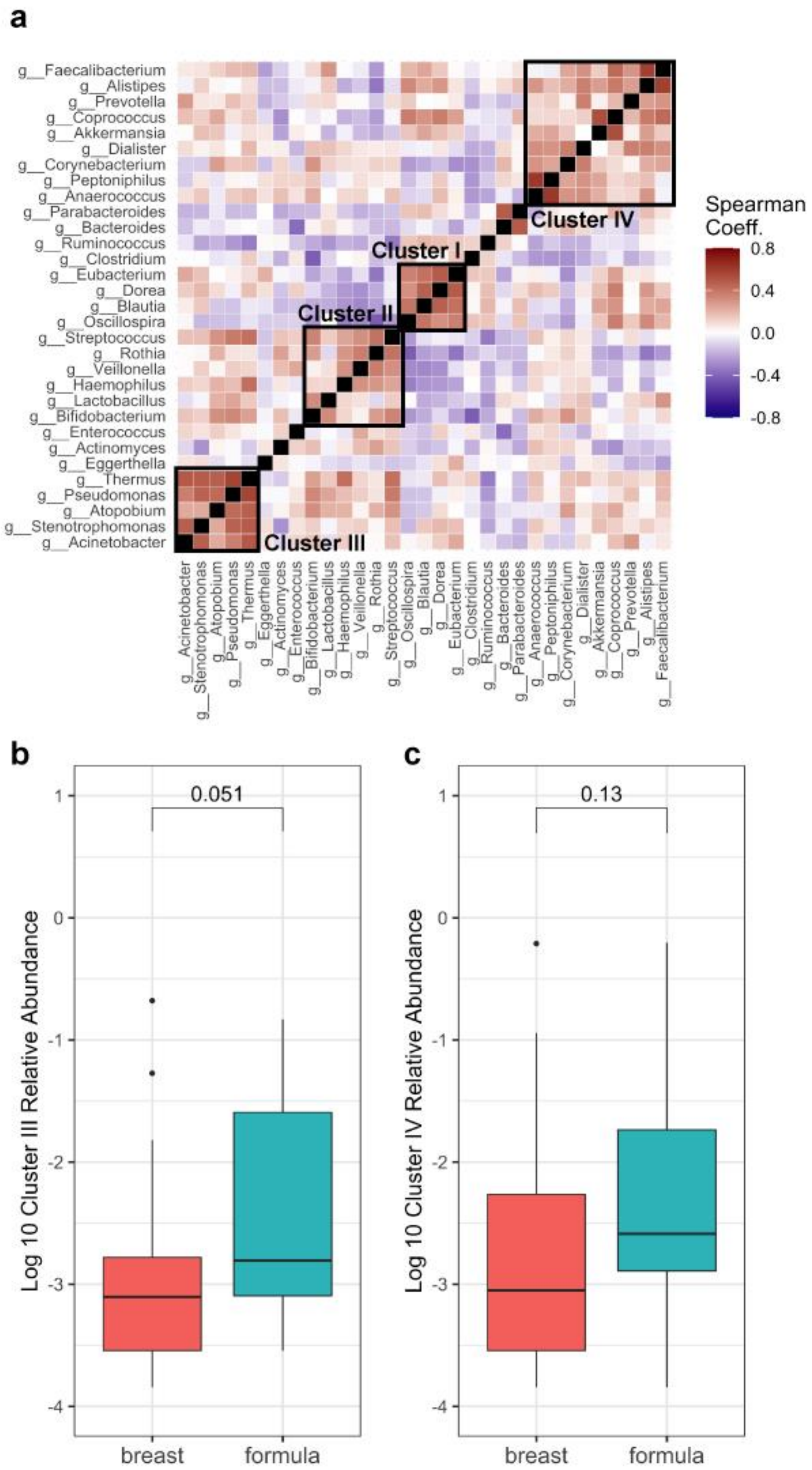
a
Spearman Coeff.
0.8
0.4
0.0
-0.4
-0.8
Cluster I
Cluster II
Cluster III
Cluster IV
g__Faecalibacterium
g__Alistipes
g__Prevotella
g__Coprococcus
g__Akkermansia
g__Dialister
g__Corynebacterium
g__Peptoniphilus
g__Anaerococcus
g__Parabacteroides
g__Bacteroides
g__Ruminococcus
g__Clostridium
g__Eubacterium
g__Dorea
g__Blautia
g__Oscillospira
g__Streptococcus
g__Rothia
g__Veillonella
g__Haemophilus
g__Lactobacillus
g__Bifidobacterium
g__Enterococcus
g__Actinomyces
g__Eggerthella
g__Thermus
g__Pseudomonas
g__Atopobium
g__Stenotrophomonas
g__Acinetobacter
b
0.051
Log 10 Cluster III Relative Abundance
1
0
-1
-2
-3
-4
breast
formula
c
0.13
Log 10 Cluster IV Relative Abundance
1
0
-1
-2
-3
-4
breast
formula

**Supplementary Fig. 4 Clusters III and IV are not differentially abundant between breastfed and formula-fed infants**. (A) Heatmap based on the covariance of bacterial genera coloured by Spearman correlation coefficient. Axes are arranged based on Spearman correlation distance and Ward linkage. (B) Comparison of cluster III and IV relative abundances across feeding groups using the Mann-Whitney U test.

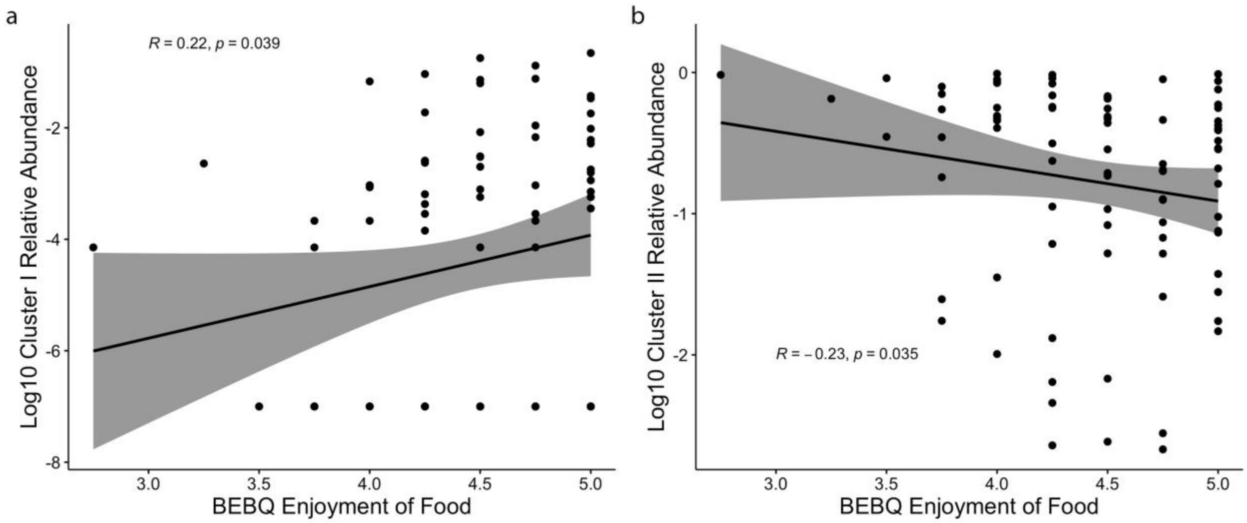


**Supplementary Fig. 5 BEBQ enjoyment of food is also significantly correlated with cluster abundances.** (A-B) Spearman correlation coefficient and significance for clusters I and II and BEBQ enjoyment of food.

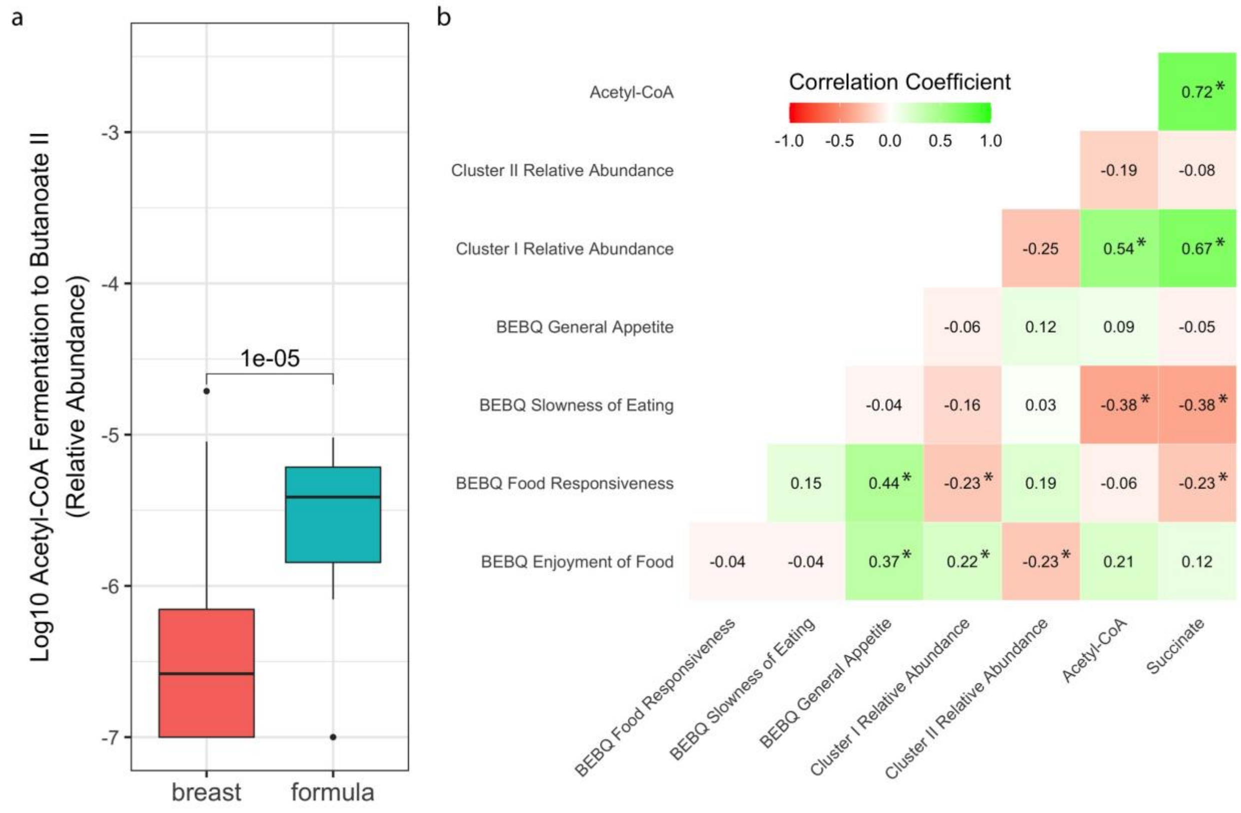


**Supplementary Fig. 6 Pathway relative abundance differs significantly between breastfed and formula-fed infants, and is significantly correlated with cluster I relative abundance.** (A) Acetyl-CoA fermentation to butanoate II is significantly more highly expressed in formula-fed infants according to the Mann-Whitney U test. (B) Heatmap of correlation matrix between cluster abundance, food responsiveness, and SCFA pathways coloured by Spearman correlation coefficient. Significance ($p < 0.05$) is marked with a star (*).
